# *Cdc42* activity in Sertoli cells is essential for maintenance of spermatogenesis

**DOI:** 10.1101/2021.04.26.441535

**Authors:** Bidur Bhandary, Anna Heinrich, Sarah J. Potter, Nancy Ratner, Tony DeFalco

**Affiliations:** Division of Reproductive Sciences, Cincinnati Children’s Hospital Medical Center, Cincinnati, OH 45229 U.S.A.; Division of Experimental Hematology and Cancer Biology, Cincinnati Children’s Hospital Medical Center, Cincinnati, OH 45229 U.S.A.; Department of Pediatrics, University of Cincinnati College of Medicine, Cincinnati, OH 45267 U.S.A.

**Keywords:** Sertoli cell, cell polarity, cell junction, Cdc42, Rho GTPase, spermatogenesis, blood-testis barrier, ectoplasmic specialization

## Abstract

Sertoli cells are highly polarized testicular supporting cells that simultaneously nurture progressively maturing germ cells. Proper localization of polarity protein complexes within Sertoli cells, including those responsible for blood-testis barrier formation, are vital for successful spermatogenesis. However, the mechanisms and developmental timing that underlie the establishment of polarity are poorly understood. To investigate this aspect of testicular function, we conditionally deleted *Cdc42*, encoding a Rho GTPase involved in regulating cell polarity, specifically in Sertoli cells. *Cdc42* deletion disrupted adult Sertoli cell maturation and localization of polarity proteins, but did not affect fetal and early postnatal testicular development, nor the onset of the first wave of spermatogenesis. By early adulthood, however, conditional knockout males exhibited a loss of spermatogenic cells, resulting in a complete lack of sperm. These findings demonstrate that *Cdc42* plays an essential role in establishing adult Sertoli cell polarity and, thus, maintaining steady-state spermatogenesis and healthy sperm production.

## INTRODUCTION

Cell polarity, in which different morphologies or molecular components are asymmetrically distributed across a cell, is a critical feature of many cell types. Asymmetries in polarized cells such as epithelial cells, migratory cells, and stem cells are required for barrier functions, directed cell movements, and cell fate decisions, respectively (Campanale et al., 2017). Within epithelial cells, including the Sertoli cells of the adult mammalian testis, asymmetry across the apical-basal axis is a key cellular characteristic (Gao and Cheng, 2016; Gao et al., 2016). Among the core molecular machinery underlying apicobasal cell polarity are 3 main protein polarity complexes: 1) the Par3/Par6/aPKC (PARD3/PARD6A/PRKCZ) complex, 2) the Crumbs (CRB3) complex, and 3) the Scribble (SCRIB) complex. The Par3/Par6/aPKC and Crumbs complexes are typically localized to apical membranes and cell junctions, while the Scribble complex is basolaterally associated (Campanale et al., 2017; Gao et al., 2016); these specific subcellular localizations are vital to maintain unique functions of different cell compartments.

Sertoli cells are the supporting epithelial cells of the testis, whose function is to nurture germ cells in their developmental path from gonocytes in the fetal gonad to spermatozoa in the adult testis. Sertoli cells are the first male-specific cells to differentiate in the fetal gonad and, in conjunction with germ cells, they undergo tubular morphogenesis to form seminiferous tubules, the structural and functional units of the testis (Cool et al., 2012). As the testis develops, Sertoli cells undergo significant changes in their cellular behavior. From fetal stages until approximately 10-14 days after birth, Sertoli cells actively divide, increasing the diameter of the seminiferous tubules and the number of potential spermatogonial stem cell (SSC) niches in the testis. At the end of this time period, however, Sertoli cells exit the cell cycle and remain quiescent throughout peri-pubertal and adult stages (Kluin et al., 1984; Orth, 1982). As they mature, Sertoli cells also undergo significant changes in the expression and localization of a number of Sertoli-specific proteins that are critical for their supportive roles in the testis. Anti-Mullerian hormone (AMH) is down-regulated in mature Sertoli cells; GATA1 is up-regulated and is expressed only in a subset of seminiferous tubules, which is dependent on feedback from specific stages of germ cells within the tubule (Yomogida et al., 1994); and androgen receptor (AR), the receptor for testosterone produced by interstitial Leydig cells, is translocated from the Sertoli cell cytoplasm to the nucleus upon maturation (Sharpe et al., 2003).

Spermatogenesis is maintained, in large part through the guidance of Sertoli cells, via a balance of mitotic and differentiating steps of SSCs, the germline stem cell population (sometimes referred to more broadly as a stem/progenitor population called undifferentiated spermatogonia). SSCs proliferate and self-renew to maintain their population, while a subset of SSCs proceed through a differentiation program, launched in response to retinoic acid, passing through differentiated spermatogonial, spermatocyte, and spermatid stages before finally producing testicular spermatozoa (Franca et al., 2016; Griswold, 2016; Kanatsu-Shinohara and Shinohara, 2013; Oatley and Brinster, 2012; Yoshida, 2010). During this process, Sertoli cells nurture all stages of germ cells as they proliferate and differentiate to give rise to sperm.

During adult steady-state spermatogenesis, Sertoli cell polarization is essential to support various stages of spermatogenic cells and to compartmentalize its functions, including maintaining germline stem/progenitor cells in the basal compartment of the seminiferous tubule while simultaneously promoting spermiogenesis in the apical compartment (Franca et al., 2016; Oatley and Brinster, 2012). A unique property of Sertoli cells that allows them to compartmentalize their various cellular interactions is a series of tight junctions that forms the blood-testis barrier (BTB), which sequesters meiotic and post-meiotic cells within the apical compartment and provides an immune-privileged niche for the latter stages of spermatogenesis. Neo-antigens from meiotic and post-meiotic cells arise in developing spermatocytes and spermatids after systemic tolerance is established, which likely renders these germ cells “foreign” by the adaptive immune system. Therefore, the physical sequestration of these antigens is a critical function of Sertoli cells, and suggests that Sertoli polarization, including proper basal localization of the BTB, is vital for spermatogenesis (Gao and Cheng, 2016; Gao et al., 2016).

Although Sertoli cell polarity is vital for successful spermatogenesis, when these cells become polarized, and through what mechanisms, is currently poorly understood. A potential player in this process is Rho GTPase signaling, which has been linked to cell polarity and other cellular processes (e.g., cell cycle, cell migration, and cytoskeletal structure) in diverse biological contexts (Etienne-Manneville and Hall, 2002; Mack and Georgiou, 2014; Ngok et al., 2014). A central member of the Rho GTPase family is *Cdc42*, which has been implicated in several aspects of cellular biology, including establishing polarity in epithelial cells (Erickson and Cerione, 2001; Kroschewski et al., 1999; Melendez et al., 2011). When activated in its GTP-bound state, CDC42 binds to and activates numerous effector proteins. CDC42 is thought to interact with Par6 of the Par3/Par6/aPKC complex (Gao et al., 2016). aPKC regulates both the BTB and the ectoplasmic specialization (ES), an actin-rich atypical adherens junction that tethers spermatids to the apical compartment of the Sertoli cell (Wong et al., 2008). *In vitro* approaches have implicated *Cdc42* in regulation of Sertoli-Sertoli BTB dynamics during the seminiferous epithelial cycle (Wong et al., 2010) and in Sertoli-germ cell interactions at the apical ES mediated by the effector protein IQGAP1 (Lui et al., 2005). However, little is known about the *in vivo* role of *Cdc42* within Sertoli cells during testicular differentiation and spermatogenesis.

Here we show, using *Cdc42* conditional knockout (cKO) mice, that *Cdc42* activity in Sertoli cells *in vivo* is not essential for fetal testicular differentiation and the postnatal onset of the first wave of spermatogenesis, but is required for latter stages of first-wave spermatogenesis beyond the round spermatid stage. *Cdc42* cKO mice exhibited severe defects in testicular structure and function beginning at peripubertal stages that continued through adulthood, including disruption of Sertoli apicobasal polarity and a loss of germ cells by the onset of sexual maturity, leading to a total lack of sperm. Overall, these findings demonstrate that *Cdc42* activity, and its role in mediating apicobasal polarity, in Sertoli cells is dispensable for early aspects of testicular development, but is critical for adult steady-state spermatogenesis.

## RESULTS

### *Cdc42* in Sertoli cells is essential for adult steady-state spermatogenesis

To analyze the role of *Cdc42* in Sertoli cells *in vivo*, *Cdc42* conditional knockout mice were generated using *Dhh*-Cre (referred to as “cKO” hereafter), which we recently confirmed is highly specific and effective for targeting Sertoli cells starting as early as E12.5 (Heinrich et al., 2020). Previously published cell-type-specific and single-cell transcriptome studies indicated that *Cdc42* is highly expressed in multiple testicular cell types, including Sertoli cells, starting at the earliest stages of testicular differentiation until adulthood (Figures S1A-S1D) (Green et al., 2018; Jameson et al., 2012; Stevant et al., 2018). Therefore, if *Cdc42* has early and/or essential roles in Sertoli cell function, *Dhh-*Cre-mediated deletion will define these early roles in Sertoli cells.

In 2-month-old (P60) adult cKO male mice testis size was severely reduced relative to heterozygous control littermates (Figure 1A). Consistent with this observation, we found that both testis weight and testis:body weight ratio in cKO males were significantly reduced (75% and 60% reductions, respectively) relative to control littermates (Figures 1B and 1C).

**Figure 1.**
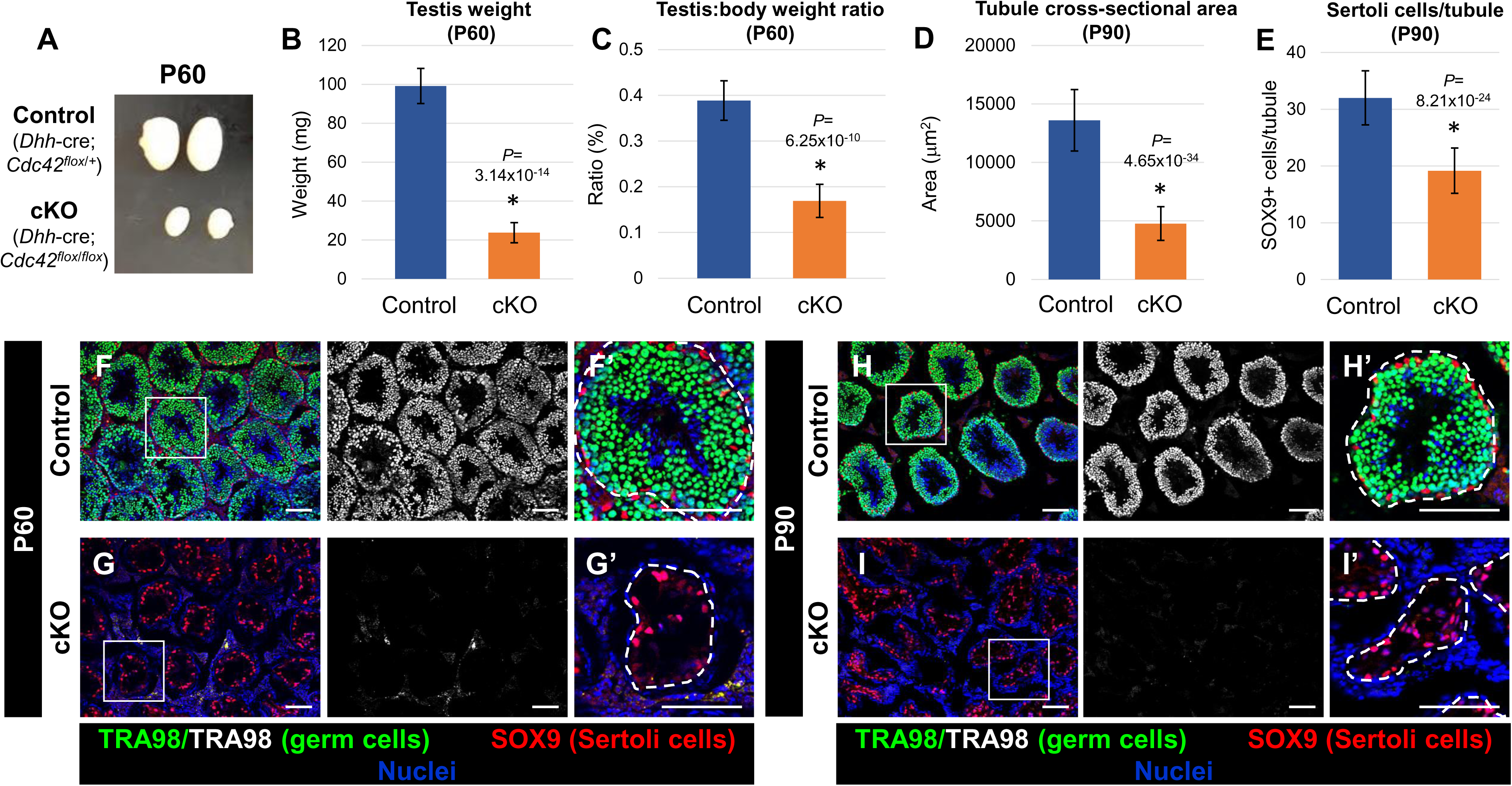
Sertoli-cell-specific ablation of *Cdc42* causes complete loss of germ cells in the adult testis. (A) Image showing adult (P60) testis size decrease in cKO (*Dhh*-Cre;*Cdc42^flox^*^/^*^flox^*) testes versus *Dhh*-Cre;*Cdc42^flox^*^/+^ heterozygous control testes. (B and C) Graphs showing average testis weight (B) and average testis weight:body weight ratio (C) of P60 control *Dhh*-Cre;*Cdc42^flox^*^/+^ males versus cKO males (n=6 control and n=4 cKO testes, each from independent males). (D and E) Graphs showing average tubule cross-sectional area (D) and average number of SOX9+ cells per tubule (E) of P90 control (*Dhh*-Cre;*Cdc42^flox^*^/+^) versus cKO tubules (n=15 tubules from each of 3 independent males). (F-I) Immunofluorescence images of P60 (F and G) and P90 (H and I) adult control *Dhh*-Cre;*Cdc42^flox^*^/+^ (F and H) and cKO (G and I) testes, showing absence of TRA98+ germ cells in cKO testes. F’-I’ are higher-magnification images of the boxed regions in F-I. Dashed lines indicate tubule boundaries. Scale bars, 100 µm. Data in B-E are shown as mean ± SD. *P* values were calculated via a two-tailed Student t-test.

To investigate the possibility that reduced cKO testis size occurred due to a decrease in germ cell number, we performed immunofluorescence for TRA98, which labels all germ cells within the seminiferous tubules. There was a complete loss of TRA98-positive cells in the lumen of seminiferous tubules of cKO mice as early as P60 (Figures 1F and 1G), which resulted in a significant reduction in cKO seminiferous tubule cross-sectional area (Figure 1D). We examined the expression of SOX9, which labels Sertoli cells, and found that adult cKO mice exhibited a significant reduction (∼40% reduction) in the number of Sertoli cells per tubule (Figures 1E-1I).

We further confirmed the loss of germ cells in cKO testes by staining for markers of germ cells at mitotic, meiotic, and post-meiotic stages, using the markers GFRA1 (undifferentiated spermatogonia), ZBTB16 (also known as PLZF; undifferentiated and differentiating spermatogonia), STRA8 (differentiating spermatogonia and preleptotene spermatocytes), γ-H2AX (spermatocytes), and H1T (pachytene spermatocytes through round spermatids) (Figures S2A-S2F). We did not detect germ cells of any spermatogenic stage in adult cKO testes. Consistent with these findings, quantitative PCR (qPCR) analyses of whole adult testes revealed that multiple germ cell genes examined, such as *Pou5f1* (also known as *Oct4*; marks undifferentiated and differentiating spermatogonia), *Stra8* (differentiating spermatogonia and preleptotene spermatocytes), and *Ddx4* (also known as *Mvh* or *Vasa*; expressed most strongly in spermatocytes and spermatids) were all significantly downregulated (100 to 1,000-fold) in mutant testes (Figure S2G).

### Lack of *Cdc42* inhibits development of adult Sertoli cells

To assess the role of *Cdc42* in Sertoli cell development, differentiation, or maturation, we examined the expression of several Sertoli-specific markers in adult control and cKO testes. GATA1, which is down-regulated by feedback from differentiated germ cells at certain stages of the spermatogenic cycle (Yomogida et al., 1994), was expressed in virtually every tubule in cKO testes, as opposed to expression in a subset of tubules observed in control testes (Figures 2A and 2B). AMH expression, which is down-regulated in adult control Sertoli cells, was also undetectable in the cKO testis via immunofluorescence or qPCR (Figures 2C and 2D; data not shown). Vimentin, expressed strongly in the basal cytoplasm near the nucleus in control Sertoli cells, was also strongly expressed in cKO Sertoli cells; however, cKO Sertoli cells displayed an aberrant subcellular localization of Vimentin in which it was expressed throughout the entire cell (Figures 2E and 2F). In contrast to control Sertoli cells, in which AR is strictly nuclear-localized, cKO Sertoli cells exhibited diffuse expression of AR in the cytoplasm, reminiscent of immature Sertoli cells (Figures 2G and 2H). We also examined the expression of GDNF, an essential factor for the maintenance of SSCs, which is expressed in multiple cell types, including Sertoli cells (Chen et al., 2014; Chen et al., 2016; Meng et al., 2000). In cKO testes we observed strong GDNF expression in Sertoli cells, similar to control testes (Figures 2I and 2J). qPCR analysis of adult testes revealed that Sertoli-specific genes *Sox9*, *Ar*, *Cldn11*, and *Gdnf* were significantly upregulated in cKO testes compared to controls; however, these results are likely at least partly due to an increase in the proportion of Sertoli cells in cKO testes after the loss of germ cells. In contrast, *Gata1*, *Fshr*, *Inhbb*, and *Pdgfa* were not significantly affected (Figure 2K). Overall, these results indicate that some aspects of Sertoli differentiation and maturation were perturbed in cKO testes.

**Figure 2.**
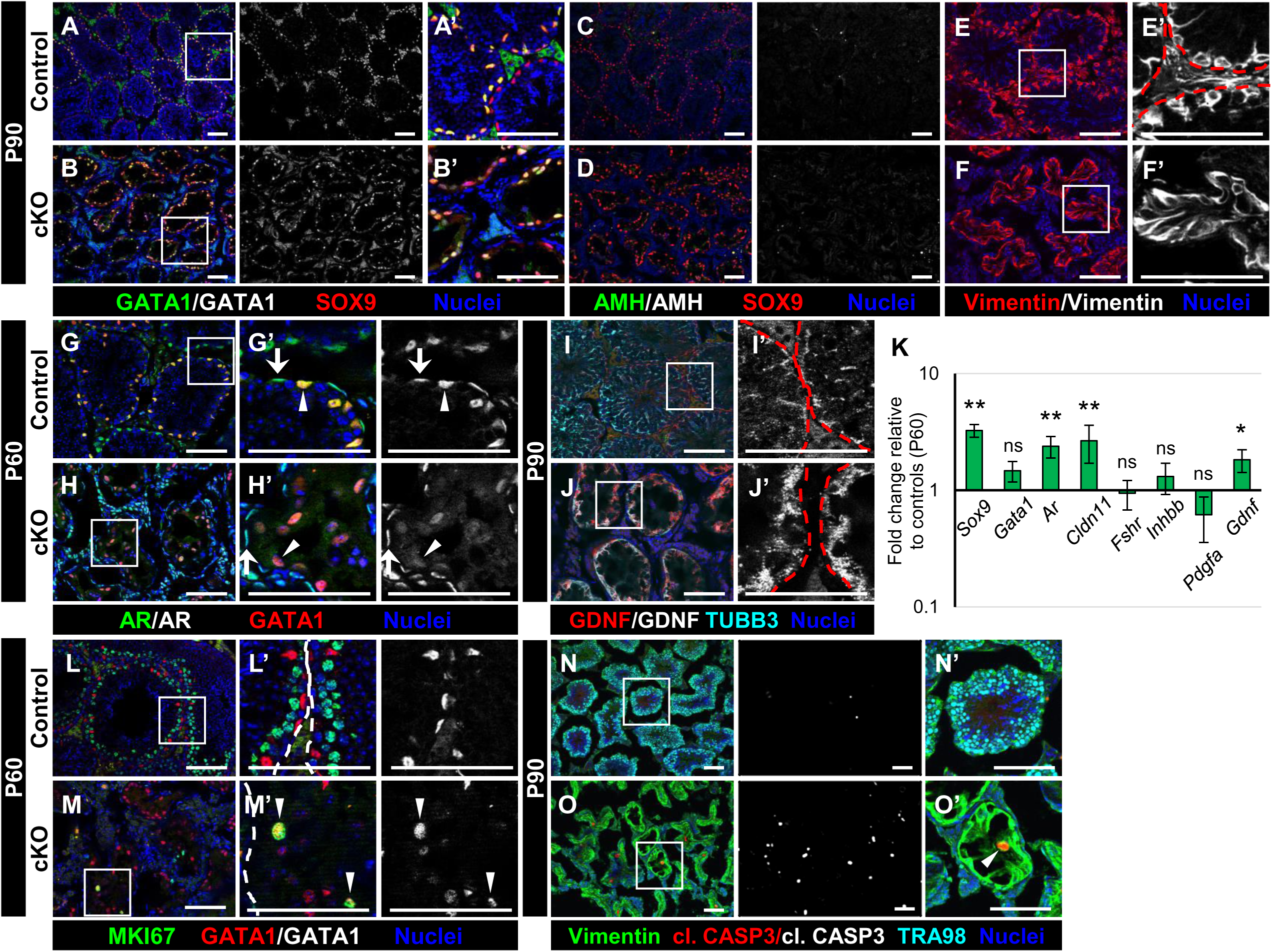
Loss of *Cdc42* disrupts development of adult Sertoli cells. (A-J,L-O) Immunofluorescence images of P90 (A-F,I-J,N-O) and P60 (G-H,L-M) adult control *Dhh*-Cre;*Cdc42^flox^*^/+^ (A,C,E,G,I,L,N) and cKO (*Dhh*-Cre;*Cdc42^flox^*^/^*^flox^*) (B,D,F,H,J,M,O) testes. A’,B’,E’-J’, and L-O’ are higher-magnification images of the boxed regions in A,B,E-J, and L-O. Dashed lines indicate tubule boundaries. (A and B) Relative to controls (A), Sertoli cells in cKO (B) testes show robust SOX9 expression but heterogeneous GATA1 expression. (C and D) Both control (C) and cKO (D) adult Sertoli cells do not express AMH. (E and F) Control (E) Sertoli cells show specific basal localization of Vimentin, but cKO (F) cells show diffuse Vimentin localization throughout the entire cell. (G and H) Control (G) testes show nuclear localization of AR in both Sertoli cells (arrowhead in G’) and peritubular myoid cells (arrow in G’), but cKO (H) cells have normal AR localization in peritubular myoid cells (arrow in H’) with diffuse cytoplasmic AR localization in Sertoli cells (arrowhead in H’). (I and J) Both control (I) and cKO (J) Sertoli cells exhibit robust GDNF expression. (K) qPCR analyses of Sertoli-expressed genes in P90 *Dhh*-Cre;*Cdc42^flox^*^/+^ control versus cKO testes (n=4 testes each for controls and cKO, all from independent males). Data in K is shown as mean fold change ± SD. **, *P*<0.01; *, *P*<0.05. *P* values were calculated via a two-tailed Student t-test. (L and M) Control (L) Sertoli cells are MKI67-negative, but cKO (M) Sertoli cells expressing MKI67 (arrowheads in M’) are occasionally observed. (N and O) CC3-positive (apoptotic) Sertoli cells are rarely detected in control (N) testes, but are often observed in cKO (O) testes (arrowhead in O’). Scale bars, 100 µm.

### *Cdc42* is dispensable for onset of postnatal Sertoli quiescence, but is required for its maintenance in the adult

*Cdc42* has been implicated in cell cycle regulation, so we examined whether *Cdc42* deletion induced changes in cell cycle status of adult Sertoli cells. By adulthood, normal Sertoli cells are mitotically quiescent, while germ cells continue to proliferate. However, in adult cKO testes, an increased number of Sertoli cells expressed MKI67 (also known as Ki67), suggesting that the Sertoli cell cycle is affected by *Cdc42* deletion (Figures 2L and 2M). We next sought to address if *Cdc42* loss affects Sertoli cell survival. We observed a significant increase in cleaved Caspase 3 (from now on referred to as CC3) staining in cKO testes; co-staining revealed that apoptotic cells were Sertoli cells (Figures 2N and 2O), consistent with the reduction in Sertoli cell number seen in cKO tubules (Figure 1E). These observations indicate that not only is *Cdc42* vital for proper development and function of Sertoli cells, it is also critical for adult Sertoli cell survival and maintenance of cell cycle arrest.

Given our results regarding increased cell cycle activity and cell death of Sertoli cells in the adult cKO testis, we assessed whether *Cdc42* is required for postnatal Sertoli cells to enter mitotic arrest and become quiescent. We examined postnatal testes at P15 and P24, stages by which Sertoli cells should have entered cell cycle arrest. We observed that Sertoli cells were all MKI67-negative at P15 and P24, apart from very rare cycling Sertoli cells, in both controls and cKO testes (Figure S3A-S3D). These data indicate that *Cdc42* is not required for the onset of quiescence in postnatal Sertoli cells.

### Disruption of germline and Sertoli cell development occurs in postnatal cKO testes

Given that germ cells were completely absent and Sertoli cells were significantly impacted at P60, we wanted to establish when these defects began to manifest during testicular development. We started by examining cKO testes at P24, a stage in which the BTB has been established for at least a week, Sertoli cells are in the process of maturation, and meiotic and post-meiotic germ cells can be found in the seminiferous epithelium.

At this stage, there were obvious Sertoli cell defects. Sertoli cell nuclei were often observed within the lumen of cKO tubules, as opposed to normal localization near the basement membrane (Figures 3A and 3B). CLDN11 was frequently diffusely expressed throughout the entirety of the tubules in cKO testes, as opposed to the basal expression outlining the BTB that was observed in controls, although in some cKO tubules there was a grossly normal CLDN11 expression pattern (Figures 3C and 3D). We observed similar disruption in the localization of AR; AR was diffusely expressed throughout cKO Sertoli cells, similar to our observations in the cKO adult (Figures 3E and 3F).

**Figure 3.**
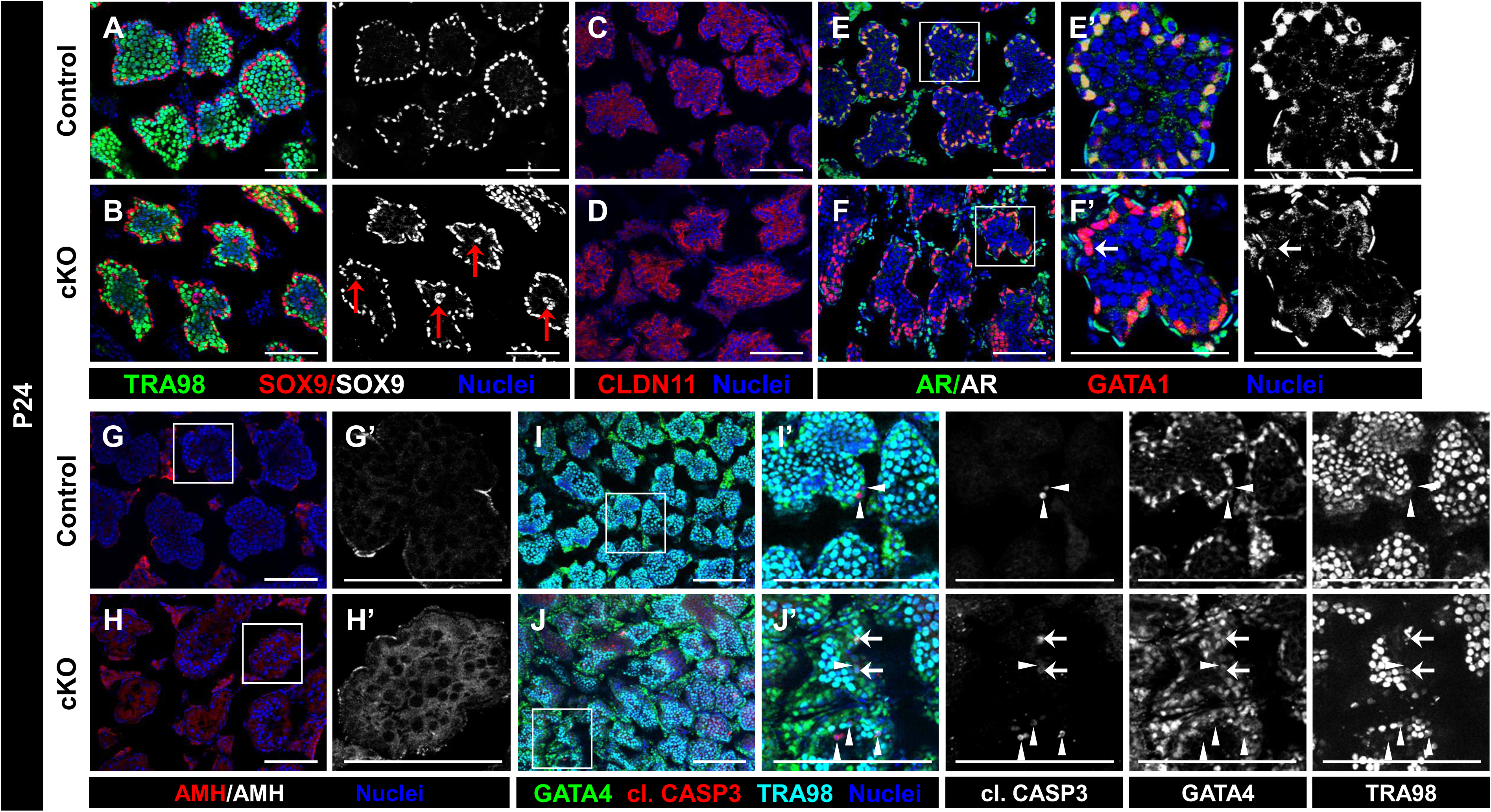
*Cdc42* deletion causes defects in differentiation and induces apoptosis of peripubertal Sertoli cells. (A-J) Immunofluorescence images of P24 control *Dhh*-Cre;*Cdc42^flox^*^/+^ (A,C,E,G,I) and cKO (*Dhh*-Cre;*Cdc42^flox^*^/^*^flox^*) (B,D,F,H,J) peripubertal testes. E’-J’ are higher-magnification images of the boxed regions in E-J. (A and B) Both P24 control (A) and cKO (B) tubules contain SOX9+ Sertoli cells, but cKO tubules display mislocalized Sertoli nuclei in the center of tubule lumens (red arrows in B). (C and D) Compared to control (C) tubules, which show a ring-like localization of CLDN11 to the BTB, cKO (D) tubules have a more diffuse pattern of CLDN11 expression. (E and F) AR localization in control (E) Sertoli cells is nuclear, but is cytoplasmic and diffuse in cKO (F) Sertoli cells (arrow in F’). (G and H) Compared to controls (G), in which AMH is undetectable within tubules, it is upregulated in cKO (H) tubules. (I and J) In contrast to controls (I), in which the few apoptotic cells are mostly germ cells (arrowheads in I’), cKO (J) tubules contain an increased number of both apoptotic germ (arrowheads in J’) and Sertoli (arrows in J’) cells. Scale bars, 100 µm.

Consistent with a disruption or delay in Sertoli cell maturation, AMH expression remained high in cKO tubules relative to controls (Figures 3G and 3H).

While there was a slight, but noticeable, decrease in TRA98+ cells and tubule diameter, germ cells were still numerous in cKO testes at this stage (Figures 3A and 3B). To test whether the reduction in germ cells was due to an increase in apoptosis, we stained for CC3 and observed an increase in the number of apoptotic cells in P24 cKO testes. Co-staining with germ and Sertoli markers revealed that apoptosis occurred mostly in germ cells, although we also found apoptotic cKO Sertoli cells, albeit less frequently than apoptotic germ cells (Figures 3I and 3J).

### *Cdc42* function is essential for maintenance of Sertoli cell polarity

We next sought to determine the effects of *Cdc42* deletion on Sertoli cell polarity *in vivo*. In control adult testes, Sertoli cell nuclei are localized basally, adjacent to the basement membrane of the tubules; however, in cKO testes we often observed Sertoli cell nuclei localized within the lumen of tubules, either as individual nuclei or in small clusters (Figures 4A and 4B).

**Figure 4.**
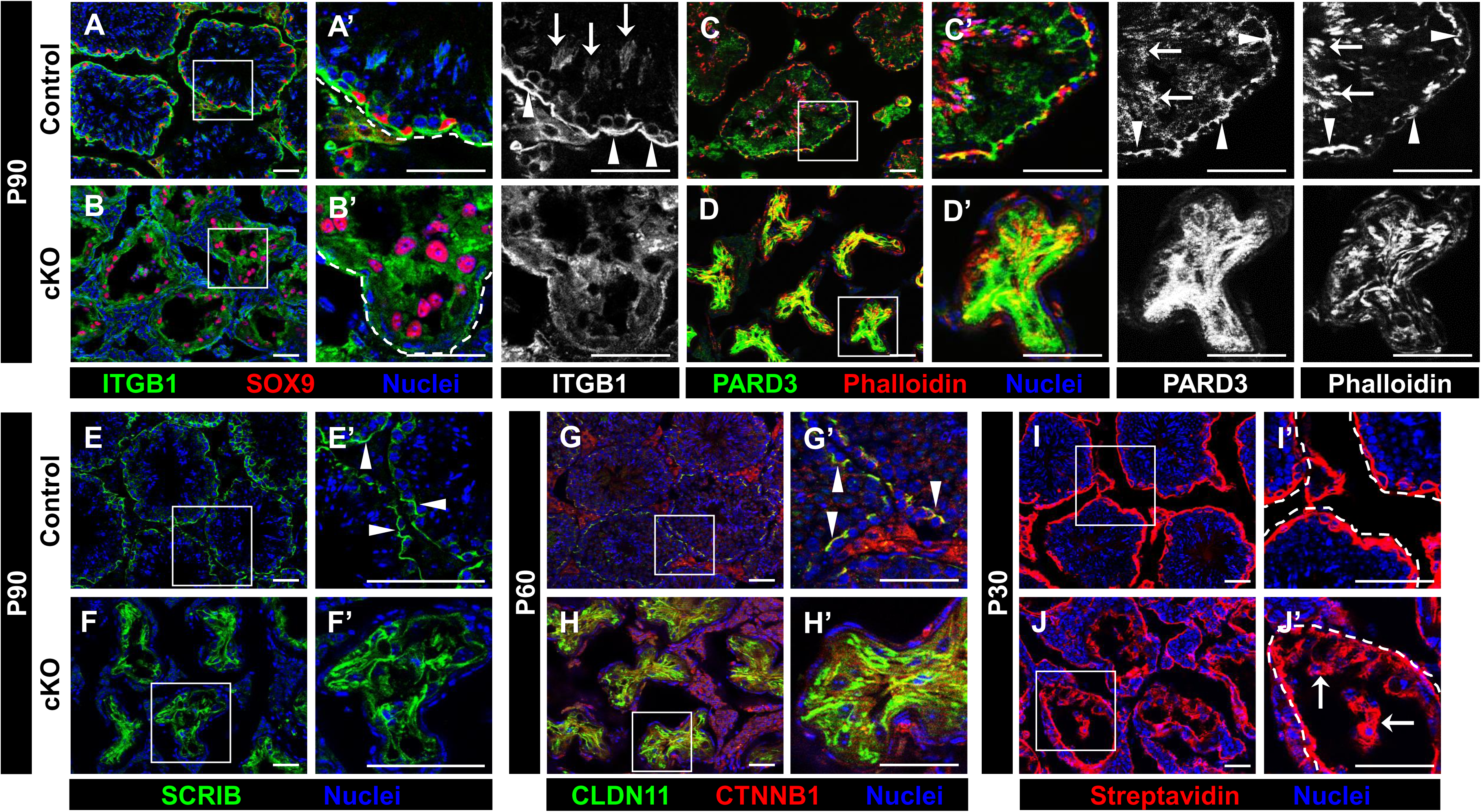
*Cdc42* activity in Sertoli cells is required for proper localization of polarity protein complexes and BTB integrity. (A-J) Immunofluorescence images of P90 (A-F), P60 (G and H), and P30 (I and J) control *Dhh*-Cre;*Cdc42^flox^*^/+^ (A,C,E,G,I) and cKO (*Dhh*-Cre;*Cdc42^flox^*^/^*^flox^*) (B,D,F,H,J) testes. A’-J’ are higher-magnification images of the boxed regions in A-J. Dashed lines indicate tubule boundaries. (A-D) In control (A and C) tubules, ITGB1, PARD3, and F-actin (via phallloidin) are enriched in the apical ES (arrows in A’ and C’) and basally in the BTB or basement membrane (arrowheads in A’ and C’), but in cKO (B and D) tubules these proteins are diffusely localized. (E and F) Control tubules (E) display specific SCRIB localization in the basal compartment (arrowheads in E’), but cKO (F) tubules exhibit diffuse localization. (G and H) In control (G) testes, CLDN11 and CTNNB1 are enriched in the BTB (arrowheads in G’), but are localized throughout entire cKO (H) tubules. (I and J) Biotin tracer injections into P30 control (I) testes reveal limited biotin presence in the interstitium and basal-most tubule cells, but in P30 cKO (J) testes there is widespread, deep penetration of biotin into the tubule lumen (arrows in J’). Scale bars, 50 µm.

Thus, having reason to suspect significant disruptions in Sertoli cell polarity, we assessed the localization of a number of polarity proteins within cKO Sertoli cells. We first examined the localization of ITGB1 (β1-integrin), an integrin protein localized to the apical portion of Sertoli cells in the ES and to the basement membrane in control testes; in contrast, in cKO testes ITGB1 was localized diffusely throughout the entire Sertoli cell (Figures 4A and 4B). A central protein of the Par polarity complex, PARD3 (Par3), was enriched in the apical compartment and at the BTB in control Sertoli cells, but in cKO cells PARD3 was distributed throughout the entire Sertoli cell (Figures 4C and 4D). We then examined the localization of F-actin via phalloidin staining, which was enriched at the actin-rich BTB and ES structures in controls, but similar to other markers it was inappropriately localized diffusely throughout cKO Sertoli cells (Figures 4C and 4D).

We next examined the expression of basally localized proteins. As expected, CLDN11 (Claudin11), CTNNB1 (β−catenin), and SCRIB (Scribble) were all localized specifically at the BTB or in the basal compartment within control tubules, but in cKO testes all were localized strongly throughout the entirety of the Sertoli cells (Figures 4E-4H).

### Barrier function of the BTB requires Sertoli *Cdc42* activity

To determine if the aberrant localization of polarity proteins correlated with changes in BTB function, we assessed its barrier function via a biotin tracer assay at P30, when the BTB should be well-established. In cKO testes, almost all tubules exhibited the presence of biotin deep within their lumen, whereas in controls biotin was only detected in the interstitium and the basal-most layer of germ cells (i.e., spermatogonia) (Figures 4I and 4J). This compromised BTB was apparent through P90, suggesting the defect at P30 was not merely a delay in BTB establishment (Figures S4A and S4B). These analyses confirm that conditional deletion of *Cdc42* results not only in disrupted polarity complexes, but in the functional disruption of the BTB.

To determine if BTB disruptions were potentially caused by a defect in testosterone production, we assessed Leydig cell development in cKO testes. We observed robust steroidogenic enzyme expression, CYP17A1 and *Cyp11a1*, in the interstitial compartment of cKO adult testes by immunofluorescence and qPCR (Figures S4C and S4D; Figure S4M). Additionally, we examined endothelial cell markers, PECAM1 and *Cdh5*, and saw grossly normal development of blood vessels in the testicular interstitium of cKO testes (Figures S4E and S4F; Figure S4M). However, we did observe ectopic expression of MKI67 in interstitial cells of cKO testes, especially within the peritubular cell layer (Figures S4G and S4H), suggesting some disruption of crosstalk between Sertoli and interstitial cells.

Given that the BTB is disrupted by *Cdc42* deletion, we tested potential consequences of this defect in terms of immune response. The BTB, via sequestration of potentially foreign germ cell neo-antigens, normally prevents the infiltration of immune cells into the lumen of the seminiferous tubules. However, in adult cKO testes there was an overall increase in testicular CD45+ immune cells, most of which were AIF1+ (IBA1+) macrophages. Macrophages were consistently observed ectopically within cKO tubules and were also increased on the periphery of the peritubular wall in cKO testes relative to controls (Figures S4I-S4L).

### *Cdc42* in Sertoli cells is essential for maintenance of peripubertal germ cells from the first wave of spermatogenesis

Although germ cells were completely absent in cKO testes by P60, germ cells were present at P24 (albeit at reduced numbers relative to controls), suggesting that the first wave of spermatogenesis initiated normally but was prematurely arrested or lost. To investigate this germ cell loss, we examined apoptosis in peripubertal testes. In P30 cKO testes there was a significant decrease in the TRA98+ cell population, accompanied by a significant increase in CC3 staining; closer inspection confirmed that the majority of apoptotic cells were germ cells (Figures 5A and 5B). In P24 cKO testes, there was only a slight decrease in TRA98+ germ cell number, but a significant number of these cells co-stained with CC3, indicating that the dramatic loss of germ cells seen at P30 begins to be manifested at this stage (Figures 5C and 5D). While we observed a significant number of CC3+ cells in cKO testes earlier in development (at P7), a similar number of apoptotic cells was seen in control testes, and the number of germ cells at this stage was indistinguishable between controls and cKO testes (Figures 5E and 5F; Figures 6A-6D). These data suggest that initial germ cell populations in the fetal and perinatal testis are unaffected by Sertoli *Cdc42* deletion and that germ cell apoptosis and loss begin during peripubertal stages.

**Figure 5.**
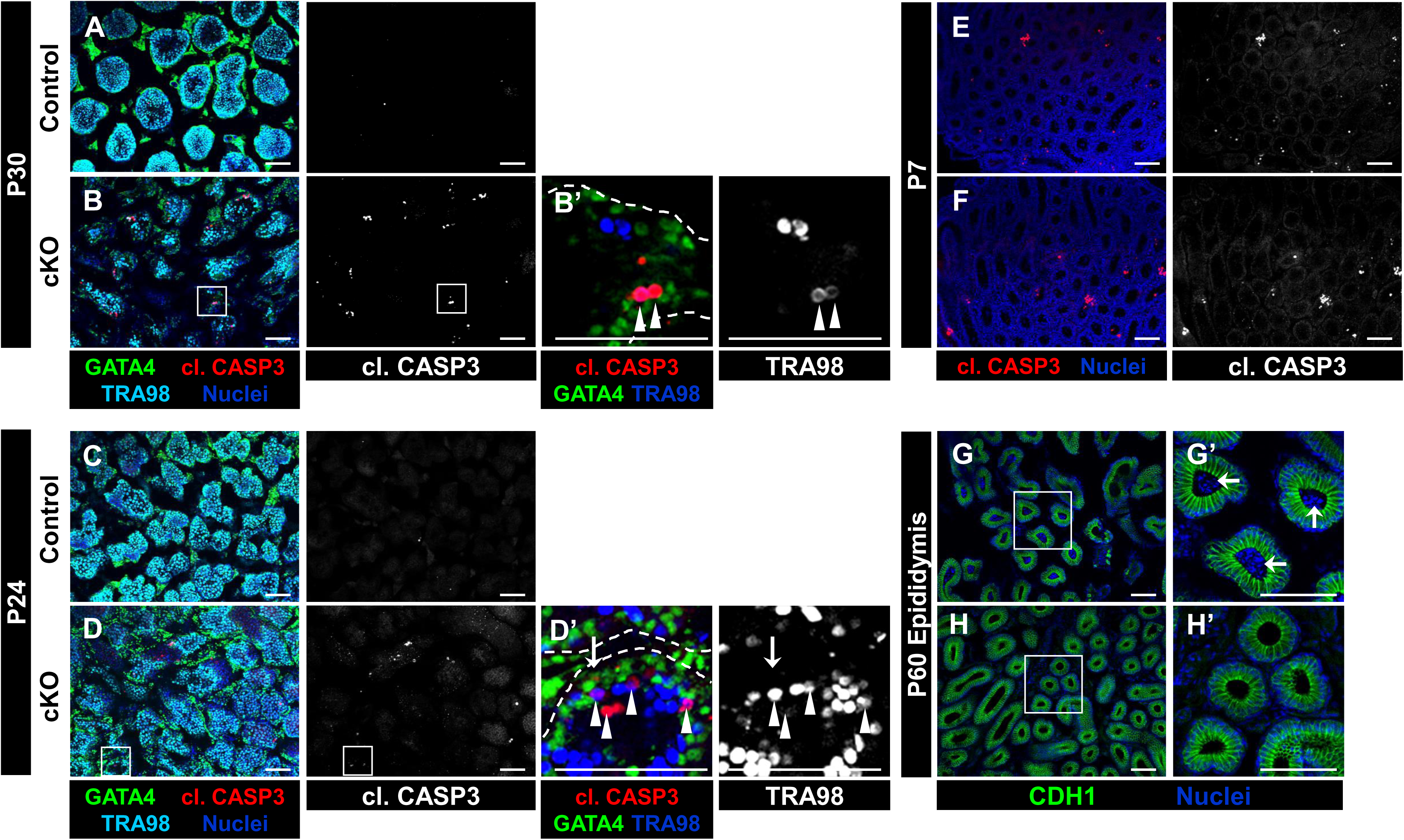
*Cdc42* cKO mice exhibit increased cell death of Sertoli and germ cells as puberty progresses, leading to a lack of epididymal sperm. (A-H) Immunofluorescence images of P30 (A,B), P24 (C,D), and P7 (E,F) testes and P60 epididymis (G,H) from control *Dhh*-Cre;*Cdc42^flox^*^/+^ (A,C,E,G) and cKO (*Dhh*-Cre;*Cdc42^flox^*^/^*^flox^*) (B,D,F,H) males. B’, D’, G’, and H’ are higher-magnification images of the boxed regions in B, D, G, and H. Dashed lines indicate tubule boundaries. (A-D) P30 control (A) and P24 control (C) testes show few CC3+ apoptotic cells, while cKO (B and D) testes show a progressive increase in apoptosis at P30 (B) relative to P24 (D), in both germ (arrowheads in B’ and D’) and Sertoli (arrow in D’) cells. (E and F) P7 control (E) and cKO (F) testes show similar numbers of apoptotic cells. (G and H) P60 control (G) epididymis contains many sperm (arrows in G’), but no sperm are detected within the cKO (H) epididymis. Scale bars, 100 µm.

**Figure 6.**
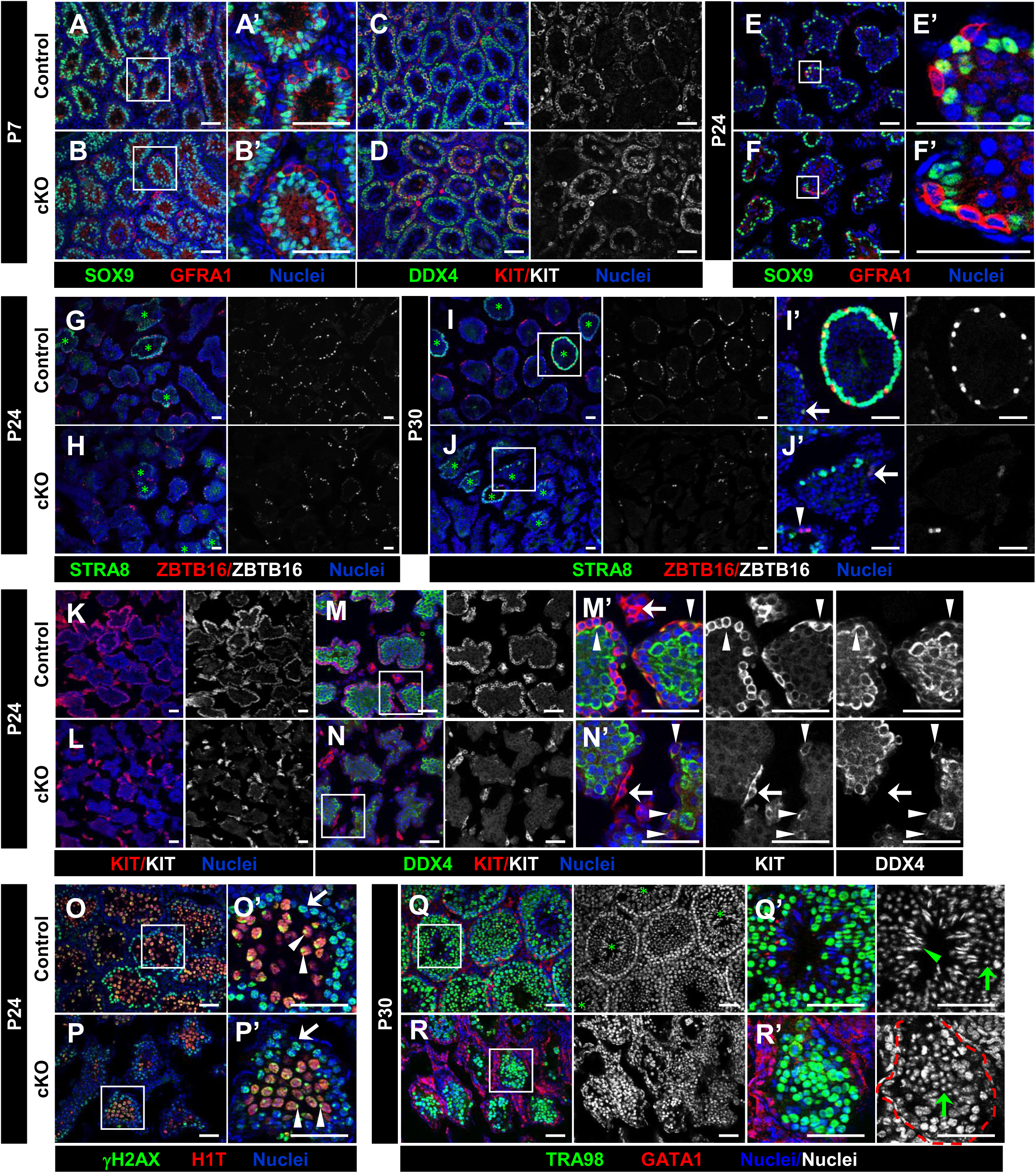
*Cdc42* cKO testes exhibit spermatogenic arrest at round spermatid stages. (A-R) Immunofluorescence images of P7 (A-D), P24 (E-H,K-P), and P30 (I,J,Q,R) postnatal control *Dhh*-Cre;*Cdc42^flox^*^/+^ (A,C,E,G,I,K,M,O,Q) and cKO (*Dhh*-Cre;*Cdc42^flox^*^/^*^flox^*) (B,D,F,H,J,L,N,P,R) testes. A’, B’, E’, F’, I’, J’, and M’-R’ are higher-magnification images of the boxed regions in A, B, E, F, I, J, and M-R. (A-D) P7 control (A and C) and cKO (B and D) testes show similar number of undifferentiated (GFRA1+) and differentiating (KIT+) spermatogonia, as well as overall germ cell number (DDX4+). (E and F) P24 control (E) and cKO (F) tubules both contain GFRA1+ cells. (G and H) Compared to P24 control (G) testes, cKO (H) testes contain fewer ZBTB16+ cells; cKO testes contain a similar number of tubules with STRA8+ cells (green asterisks in G-J indicate tubules containing STRA8+ spermatocytes), but fewer STRA8+ cells within those tubules as compared to controls. (I and J) Compared to P30 controls (I), cKO (J) testes contain considerably fewer ZBTB16+ and STRA8+ cells (arrowheads in I’ and J’ indicate ZBTB16-bright/STRA8- undifferentiated spermatogonia; arrows in I’ and J’ indicate ZBTB16-dim/STRA8+ differentiating spermatogonia). (K-N) Relative to P24 control (K and M) tubules, cKO (L and N) tubules contain dramatically fewer KIT+ differentiating spermatogonia (arrowheads in M’ and N’), but still contained KIT+ Leydig cells (arrows in M’ and N’). (O and P) Both P24 control (O) and cKO (P) testes contain γH2AX-positive (arrows in O’ and P’) and H1T-positive spermatocytes with cells XY- body-enriched γH2AX staining (arrowheads in O’ and P’), though at reduced numbers in cKO tubules. (Q and R) P30 control (Q) and cKO (R) tubules both contain round spermatids (green arrows in Q’ and R’), but elongating spermatids are only observed in control testes (green arrowhead in Q’). Scale bars, 50 µm.

To confirm that germ cells from the first wave of spermatogenesis did not give rise to spermatozoa that were released from the testis on an ongoing basis, we examined the epididymis from young adult (P60) cKO males. Whereas control males had detectable sperm within the lumen of the epididymis at P60, there were no sperm observed in cKO epididymis (Figures 5G and 5H), demonstrating that the first wave of spermatogenesis did not give rise to any sperm and suggesting that spermatogenesis was prematurely arrested or lost in the cKO testis.

### Sertoli *Cdc42* function is required for progression past the round spermatid stage in spermatogenesis

Our findings suggested that spermatogenesis was disrupted in cKO testes; therefore, we next sought to determine which spermatogenic stages were impacted by *Cdc42* conditional deletion. Consistent with what we observed via CC3 staining, early postnatal spermatogenesis at P7 was largely unaffected in cKO testes when compared to controls. Undifferentiated spermatogonia, as visualized via GFRA1 staining, were present in both control and cKO testes and all exhibited proper basal localization within the tubules (Figures 6A and 6B). Using KIT staining in conjunction DDX4 and TRA98 staining, we also determined that the differentiating spermatogonial population and overall germ cell numbers, respectively, were similarly indistinguishable between P7 control and cKO tubules (Figures 6C and 6D; Figures S5A and S5B). As for Sertoli cells, we found that P7 cKO Sertoli cells generally had similar GATA1, AMH, and AR expression as compared to controls (although AMH expression was decreased in some cKO tubules), and overall apoptosis levels of Sertoli and germ cells were similar in control versus cKO testes (Figures S5C-S5H); additionally, we saw no differences in vasculature, Leydig cells, or immune cells in cKO testes versus controls (Figures S6A-S6F). These data overall indicate that early postnatal testis differentiation and the initiation of the first wave of spermatogenesis was unaffected in cKO testes.

Germ cell loss began to increase by P24 in cKO testes, so we next examined different germ cell types at that age to determine any changes in spermatogenesis. GFRA1+ undifferentiated spermatogonia were still observed in their basal location in cKO tubules similarly to controls (Figures 6E and 6F). In contrast, assessment of ZBTB16 expression, which labels both undifferentiated (bright staining) and differentiating (dim staining) spermatogonia, revealed a reduction of these spermatogonial populations (Figures 6G and 6H); however, their basal localization did not seem to be impacted. Although there was a reduction in the number of STRA8+ cells (differentiating spermatogonia and preleptotene spermatocytes) within cKO tubules relative to controls, the number/percent of tubules containing STRA8+ preleptotene spermatocytes was not affected (Figures 6G and 6H). Overall, these data indicate that there was a reduction in the number of both undifferentiated and differentiated cells in cKO testes at P24.

Consistent with our observations of increased apoptosis and germ cell loss by P30 in cKO testes, at this stage we also observed some signs of disrupted spermatogenesis. Both ZBTB16+ undifferentiated spermatogonia and STRA8+ differentiating spermatogonia and preleptotene spermatocytes were severely reduced in cKO tubules when compared to controls. Interestingly, while the number of STRA8+ cells per tubule was significantly reduced, the number of tubules containing STRA8+ cells was indistinguishable between control and cKO testes (Figures 6I and 6J), similar to what we observed at P24. Given the loss of undifferentiated spermatogonia that we observed in cKO testes, we assessed the differentiating spermatogonial pool via KIT expression, and found a dramatic decrease in differentiating spermatogonia at P24 (Figures 6K-6N). We also found a decrease in spermatocyte and round spermatid populations at P24, as marked by γ-H2AX (official name H2A.X variant histone) and H1T (official name H1F6); however, morphology of these cell types was unaffected, including the enrichment of γ- H2AX in the XY body of pachytene spermatocytes and a condensed chromocenter within round spermatids (Figures 6O and 6P). While we were able to confirm the presence of round spermatids via H1T staining and nuclear morphology in both control and cKO tubules, elongating spermatids were never observed in any P30 tubules, in contrast to control animals (Figures 6Q and 6R), indicating that spermatogenesis is arrested at the round spermatid stage in cKO males.

### *Cdc42* expression in Sertoli cells is dispensable for fetal testis differentiation

To understand the role of Sertoli-cell-specific *Cdc42* function in fetal testis differentiation and development, we examined several parameters of Sertoli and germ cells in E18.5 fetal testes. Our analyses revealed no differences in general tubule size, structure, and marker expression in major cell types within cKO testes, including Sertoli cells, germ cells, and Leydig cells. In cKO fetal testes, AMH- and SOX9-positive Sertoli cells were arranged normally along the basal surface of the seminiferous tubule, with similar expression of CTNNB1 as compared to control tubules (Figures 7A-7D). Sertoli cells in both cKO and controls were in active cell cycle, as defined by MKI67 expression, and germ cells were centrally localized within the tubule and were in cell cycle arrest (MKI67-negative) (Figures 7E-7H). Leydig cell development and male-specific vascularization of the testis were also normal in cKO samples (Figures 7I-7J). Finally, we observed no significant difference in CD45+ cells, which were mostly located outside the seminiferous tubule lumen, in control versus cKO testes (Figures 7K-7L). Overall, our results demonstrate that *Cdc42* in Sertoli cells is not essential for fetal testicular somatic and germ cell development.

**Figure 7.**
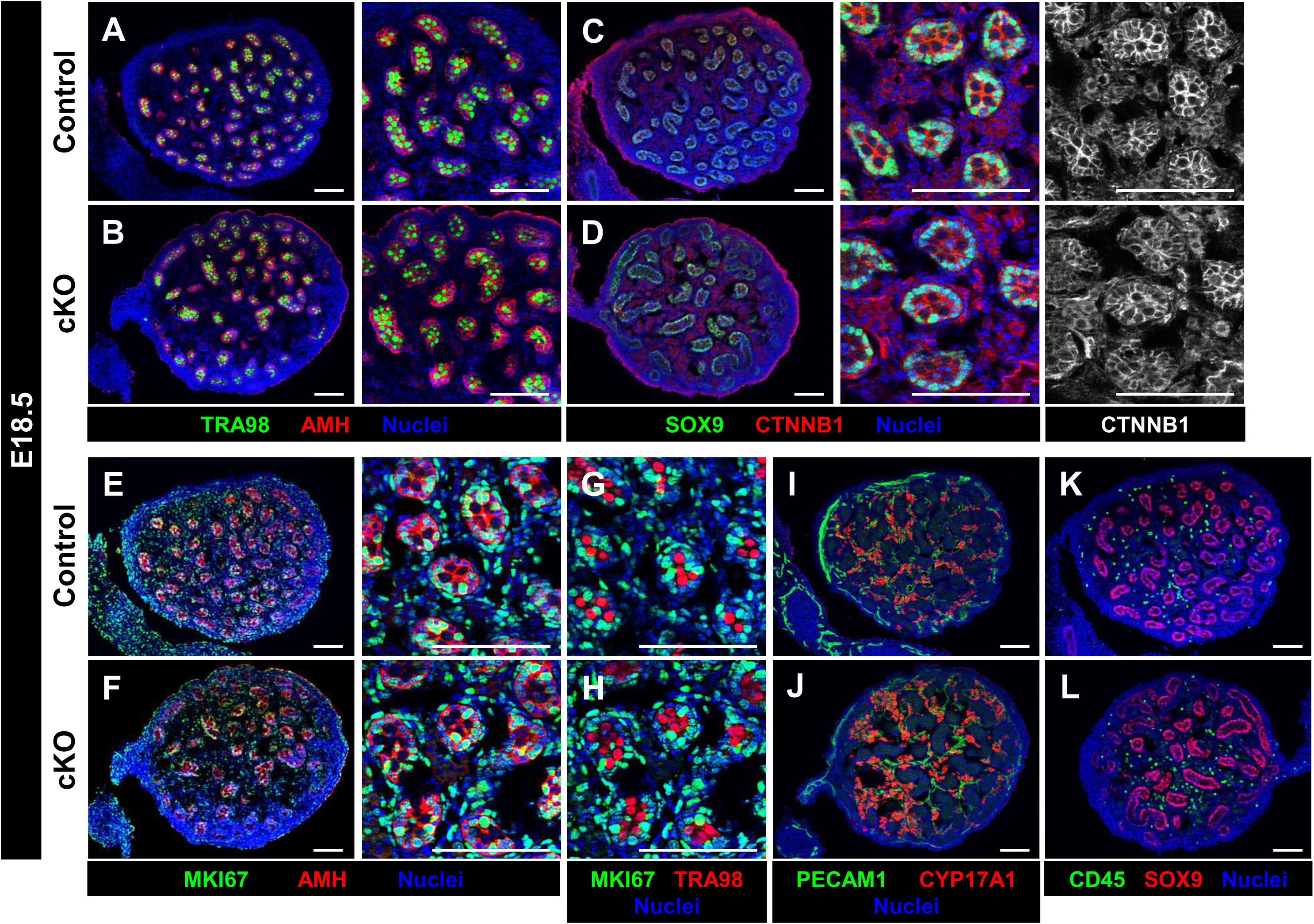
*Cdc42* in Sertoli cells is dispensable for fetal testicular differentiation and development. (A-L) Immunofluorescence images of E18.5 control *Dhh*-Cre;*Cdc42^flox^*^/+^ (A,C,E,G,I,K) and cKO (*Dhh*-Cre;*Cdc42^flox^*^/^*^flox^*) (B,D,F,H,J,L) testes. (A-D) Control (A and C) and cKO (B and D) testes contain similar numbers of germ (TRA98+) and Sertoli (AMH+/SOX9+) cells, and display similar expression of CTNNB1 within tubules. (E and F) Both control (E) and cKO (F) Sertoli cells are in active cell cycle (MKI67+). (G and H) Both control (G and cKO (H) germ cells are quiescent (MKI67-negative). (I-L) Control (I and K) and cKO (J and L) testes show similar numbers of vascular endothelial (PECAM1+), Leydig (CYP17A1+), and immune (CD45+) cells. Scale bars, 100 µm.

## DISCUSSION

The Sertoli cell is a dramatic example of a polarized epithelial cell, both in its structure and in its exquisite regulation of functions that require apicobasal polarity. Spermiation, the release of sperm from the Sertoli cell into the seminiferous tubule lumen, takes place simultaneously with remodeling of the BTB that allows preleptotene spermatocytes to traverse the BTB; at the same time, germ cells enter the first stage of meiosis. These three events occur at opposite ends of the Sertoli cell and underscore the importance of apicobasal polarity, both intracellularly and at the plasma membrane. While some progress has been made in the last two decades regarding the mechanisms that underlie Sertoli cell polarity (reviewed in (Gao and Cheng, 2016; Gao et al., 2016; Wong and Cheng, 2009)), many aspects of this process are still poorly understood. In particular, the Rho GTPase *Cdc42* has been proposed to be critical for BTB dynamics, mostly using *in vitro* cell culture and dominant-negative models (Wong et al., 2010), but its role in spermatogenesis and testis differentiation *in vivo* was unclear. Here we provide *in vivo* evidence that Sertoli cells require *Cdc42* to establish apicobasal polarity critical for supporting adult steady-state spermatogenesis.

Our conditional ablation of *Cdc42* in Sertoli cells resulted in severe disruption of adult testicular structure and function, leading to a complete loss of germ cells by 2 months of age. Underlying this loss of the germline, the structure of adult cKO seminiferous tubules was significantly disrupted, with reduced Sertoli cell number and tubule diameter, likely due to increased apoptosis of Sertoli cells. Perhaps most striking was a total mislocalization of polarity protein complexes, such as the Par6/Par3/aPKC complex and the Scribble complex, and of factors localized to BTB tight junctions and ES adherens junctions, such as CLDN11, F-actin, ITGB1, and CTNNB1. These results show that *Cdc42* is an essential factor whose deletion causes severe ripple effects for the polarity and function of Sertoli cells.

Our analyses revealed a complete loss of germ cells in cKO testes, indicating defective maintenance of stem/progenitor cells in the SSC niche, leading to an exhaustion of spermatogenesis. As studies of neonatal hypothyroidism in mice revealed that Sertoli cell number is the rate-limiting aspect of SSC niche establishment and sperm production (Hess et al., 1993; Joyce et al., 1993; Oatley et al., 2011), our data is consistent with the idea that disruption of Sertoli cells, such as by disturbing apicobasal polarity, disrupts SSC maintenance.

Loss of polarity in Sertoli cells, however, may not completely account for the phenotype observed in cKO testes. *Kit* mutation causes extensive loss of germ cells (except for undifferentiated spermatogonia) and also causes a compromise in BTB function (Li et al., 2018); yet, SSCs can be maintained in this environment, indicating that BTB function is not absolutely necessary for a functional SSC niche. Recent analyses of a Sertoli-specific conditional deletion of *Rac1*, encoding another Rho GTPase, revealed BTB disruption but, interestingly, sustained maintenance of germline stem/progenitor cells; this difference may be due to less severe disruption of apicobasal polarity in *Rac1* conditional mutants as compared to *Cdc42* (Heinrich et al., 2020). However, the germ-cell-less cKO testes observed here exhibit a more severe phenotype than merely a disrupted BTB or ES. Our analyses here revealed normal expression of *Gdnf*, which is required for SSC maintenance (Meng et al., 2000), in cKO Sertoli cells, suggesting that a lack of this critical ligand is not the underlying cause of SSC loss. We propose that as-yet-unidentified effects of *Cdc42* cause the *Cdc42* cKO SSC phenotype.

There are several ways in which GTP-active CDC42 could potentially impact the cellular biology and function of Sertoli cells. The GTP-activated form of CDC42 is proposed to directly interact with PAR6DA (Par6) in the Par6/Par3/aPKC complex (Joberty et al., 2000; Lin et al., 2000), the latter of which coordinates with the Crumbs complex to ensure apical ES and BTB remodeling during spermatogenesis (Wong et al., 2008). CDC42-PAR6DA binding is thought to increase kinase activity by aPKC, which recruits the Crumbs and Scribble complexes to epithelial tight junctions, including those of the BTB (Yamanaka et al., 2001). aPKC likely has additional phosphorylation targets which may impact Sertoli cell function downstream of the Par complex, such as the junction adhesion molecules (e.g., JAM-C/Jam3) and c-src tyrosine kinase (CSK) that regulate ES Sertoli-spermatid adhesion and BTB integrity (Wong et al., 2008). Therefore, CDC42 and its downstream effectors are biologically well-situated to be an important component of establishing the Sertoli cell identity.

In addition to its roles in cell adhesion and junctional complexes, the GTP-activated form of CDC42 mediates responses to cell signaling pathways, such as TNF-α or TGF-β, that regulate BTB dynamics. It has been proposed that CDC42 (through its interactions with TGF-β or other extracellular pathways) ultimately signals directly through JNK or indirectly through Ras/ERK and p38 to elicit cellular responses (Lui et al., 2003; Wong et al., 2005; Xia and Cheng, 2005). Recently, CDC42 was implicated in mediating mTORC1 signaling downstream of NC1 peptide–mediated BTB remodeling via actin and microtubule cytoskeletal reorganization (Su and Cheng, 2019), as well as being a downstream mediator of coxsackievirus and adenovirus receptor (CXADR) signaling to regulate BTB and apical ES integrity and function (Huang et al., 2019). *In vitro* studies using dominant-negative constructs demonstrated that *Cdc42* is required for TGF-β-induced BTB breakdown (Wong et al., 2010), which normally occurs when preleptotene spermatocytes traverse the BTB, suggesting that this activity is mediated via the GTP-active form of CDC42. Further analyses of these pathways and their targets will elucidate the specific cellular effects of CDC42’s transduction of extracellular signals.

The defects observed in the cKO testis could also result from one or more of the following impacts of CDC42 within cells: changes in protein trafficking, actin cytoskeletal dynamics, cell adhesion, and/or generation of cellular processes (such as filopodia). All of these biological processes are, to a certain degree, inextricably linked to each other. In epithelial cells, the polarized localization of apical versus basal components on opposite sides of tight junctions is the sum of multiple protein trafficking events, including endocytosis, endosome recycling, and transcytosis (reviewed in (Duffield et al., 2008)). These events are often linked with the formation and maintenance of adherens junctions. In several *Drosophila* epithelia, *Cdc42* regulates the actin-mediated endocytosis of apical proteins and adherens junctions components and their passage through the endosomal pathway (Georgiou et al., 2008; Harris and Tepass, 2008; Leibfried et al., 2008). Consistent with these endocytic roles in other systems, *Cdc42* has been linked to regulating endocytosis in Sertoli cells (Wong et al., 2008, 2010; Wong et al., 2009), in particular to how cell adhesion molecules such as N-cadherin and junctional adhesion molecule-A (JAM-A/F11R) localize at the BTB (Wong et al., 2009). Differential endocytosis may be a mechanism by which polarity proteins regulate cell adhesion, which is required to maintain spermatids attached to the apical ES and which must be switched off during spermiation. While our data in this study do not support a global adhesion defect in cKO Sertoli cells, since germ cells from the first wave of spermatogenesis are maintained in the testis until puberty, the loss of elongating spermatids in cKO testes from the first wave is entirely consistent with the loss of the adherens junctions in the apical ES that are required to maintain those cells in the seminiferous epithelium.

In cKO testes, we observed extensive immune cell infiltration into the seminiferous tubules, indicative of a disrupted BTB. There are additional aspects of immune function that rely on Sertoli functions, such as the secretion of immunosuppressive molecules (reviewed in (Meinhardt and Hedger, 2011; Zhao et al., 2014)), which are worth considering in the analysis of the *Cdc42* cKO phenotype. A recently observed Sertoli cell immune function that may be linked to protein trafficking is the retrograde transcytosis of selective neo-antigens from germ cells phagocytosed by Sertoli cells into the interstitium, which informs the adaptive immune system and induces systemic tolerance (Tung et al., 2017). Therefore, defective protein trafficking may be linked to critical functions of Sertoli cells in mediating the unique immune environment of the testis that permits spermatogenesis. Another potential factor at play, as previously mentioned, is that a significant loss of germ cells may contribute to a reduction in Sertoli cell BTB barrier function, leading to an increased testicular immune response (Li et al., 2018).

Actin cytoskeletal dynamics are another potential factor to consider in Sertoli cell biology, as the actin cytoskeleton is not only vital for protein vesicular transport, but also for filipodia formation and other cellular processes. Since Sertoli cells contact multiple germ cells simultaneously, Sertoli cell cytoarchitecture specifically accommodates physical interactions with germ cells to maintain proper adhesion and intercellular signaling; the intermingling of germ and somatic cells has been carefully described in *Drosophila* gonads, and Rho GTPases also have been implicated in this phenomenon (Jenkins et al., 2003; Sarkar et al., 2007). *Cdc42* may be involved in remodeling the actin cytoskeleton to allow its unique, complex architecture to accommodate numerous germ cells at different stages of spermatogenesis.

In addition to the disrupted polarity seen in cKO Sertoli cells, we observed prolonged Sertoli cell immaturity. While AR transitions from a cytoplasmic to nuclear localization under normal conditions, AR failed to properly translocate to the nucleus in *Cdc42* cKO Sertoli cells. Although it is likely that the disrupted polarity observed in cKO cells negatively impacts spermatogenesis, this disruption in maturation cannot be ruled out as a contributor. AR function in Sertoli cells, for example, has been previously shown to be essential for the maintenance of spermatogenesis, such that Sertoli-cell-specific AR cKO animals undergo spermatogenic arrest during meiotic stages (Chang et al., 2004; De Gendt et al., 2004). However, we regularly observe round spermatids from the first wave of spermatogenesis within cKO testes; therefore, androgen signaling disruptions cannot solely account for the cKO phenotype. On the other hand, we observe a delay in down-regulation of AMH in postnatal testes and heterogeneous expression of GATA1 in adult testes, suggesting that, similar to AR mislocalization, there are defects in Sertoli cell maturation in cKO males.

Although we observed various detrimental effects of *Cdc42* cKO in adult testes, surprisingly, we observed relatively normal testicular structure and function in early postnatal and fetal cKO testis development. At E18.5 and P7, we did not observe any major impacts of *Cdc42* conditional deletion on germ or Sertoli cell development, indicating that *Cdc42*’s role in cell polarity or other processes is not required for fetal testis differentiation or early germ cell behavior in fetal or perinatal stages. These findings are consistent with our recent observations that cell polarity, as visualized by localization of polarity complexes, is not yet observed in fetal stages; additionally, we have previously shown that Sertoli-specific conditional deletion of *Rac1*, another Rho GTPase, also disrupted cell polarity in the adult testis but did not impact fetal testis development (Heinrich et al., 2020).

In summary, here we have found that *Cdc42* in Sertoli cells is absolutely essential for adult testicular function, particularly through the maintenance of apicobasal cell polarity and establishment of the BTB and apical ES within the seminiferous epithelium. The perturbation of these processes regulated by *Cdc42* ultimately affected steady-state spermatogenesis beginning at peripubertal stages and resulted in a complete disruption of germ cell maintenance by adulthood. This study provides new insights into the mechanisms underlying the establishment of Sertoli cell polarity and its critical role in ensuring spermatogenesis and fertility.

## ACKNOWLEDGMENTS

We thank C. Moon for animal husbandry; D. Meijer and Y. Zheng for providing mice; and M. Handel for H1T antibody. We also thank B. Waller, C. Spinner, and M. White for preliminary experiments. This work was supported by Cincinnati Children’s Hospital Medical Center developmental funds and internal funding, and by National Institutes of Health grants R35GM119458 and R01HD094698 to T. DeFalco.

## AUTHOR CONTRIBUTIONS

Conceptualization, B.B., S.J.P., N.R. and T.D.; Methodology, B.B., A.H., S.J.P. and T.D.; Investigation, B.B., A.H., S.J.P., L.G. and T.D.; Resources: N.R.; Writing – Original Draft, B.B., A.H., and T.D.; Writing – Review & Editing, B.B., A.H., S.J.P., N.R. and T.D.; Supervision, T.D.; Project Administration: T.D.; Funding Acquisition, T.D.

## DECLARATION OF INTERESTS

The authors declare no competing interests.

## METHODS

### Mice

All mice used in this study were housed in the Cincinnati Children’s Hospital Medical Center’s animal care facility, in compliance with institutional and National Institutes of Health guidelines. Institutional ethical approval through the Institutional Animal Care and Use Committee (IACUC) of Cincinnati Children’s Hospital Medical Center was obtained for all animal experiments. Mice were housed in a 12 hour light/12 dark cycle and had access to autoclaved rodent Lab Diet 5010 (Purina, St. Louis, MO, USA) and ultraviolet light-sterilized RO/DI constant circulation water *ad libitum*. *Dhh*-Cre (Tg(Dhh-cre)1Mejr) mice (Jaegle et al., 2003; Lindeboom et al., 2003) were obtained from Dies Meijer (University of Edinburgh) and *Cdc42^flox^* (*Cdc42^tm1Yizh^*) mice (Chen et al., 2006; Guo et al., 2013) were obtained from Yi Zheng (Cincinnati Children’s Hospital Medical Center). Both alleles were maintained on a C57BL/6J background. Homozygous *Cdc42^flox/flox^* females and hemizygous *Dhh*-Cre; heterozygous *Cdc42^flox/+^* males were bred, and their offspring were genotyped and used for analysis. *Dhh*-Cre;*Cdc42^flox/flox^* mutant mice begin to exhibit partial hind limb paralysis around 3 weeks but can live a normal lifespan if continuously provided with food on the floor of their cages after weaning (Guo et al., 2013). *Dhh-*Cre mice were genotyped using primers sense: 5’ – ACCCTGTTACGTATAGCCGA-3’ and anti-sense: 5’-CTCCGGTATTGAAACTCCAG-3’; Cre presence was confirmed via production of a 400-bp band. *Cdc42* floxed and wild-type alleles were genotyped using primers sense: 5’-TACAGTTGGTACATATTCCGATGGG-3’ and anti-sense: AGACAAAACAAGGTCCAGAAAC-3’; *Cdc2 floxed* allele was confirmed via presence of a 465-bp band and wild-type allele was confirmed via presence of a 361-bp band. For fetal time points, timed matings were arranged. Noon on the day a vaginal plug was observed was designated as E0.5.

### Immunofluorescence

Testes were dissected in PBS and fixed overnight in 4% paraformaldehyde (PFA) with 0.1% Triton X-100 (PBTx). For adult testes, the capsule was superficially punctured 10-12 times with a 27-gauge needle prior to fixation to ensure penetration of PFA. Following overnight fixation at 4°C, testes were cut in half transversely and placed again in 4% PFA for an additional 2 hours. After several washes in PBTx, fixed testis samples were processed through a sucrose:PBS gradient (10%, 15%, and 20% sucrose) before rocking at 4°C overnight in a 1:1 mixture of 20% sucrose and Optical Cutting Temperature (OCT) medium (Fischer Healthcare, TX; catalog #4585). The following day, testes samples were embedded in OCT medium and stored at −80°C until cryosectioned.

After cryosectioning at 18-20 μM, samples were washed several times in PBTx, then incubated in blocking solution (PBTx + 10% FBS + 10% bovine serum albumin [BSA]) for 1 hour at room temperature. Primary antibodies were diluted in blocking solution and applied to samples overnight at 4°C. After several washes in PBTx, fluorescent secondary antibodies (Alexa-conjugated; from Molecular Probes/Thermo Fisher, all at 1:500 dilution) and nuclear dye (2 μg/ml Hoechst 33342, catalog #H1399, Molecular Probes/Life Technologies/Thermo Fisher) were applied for 1 hour at room temperature. Following several washes in PBTx, samples were mounted on slides in Fluoromount-G (Southern Biotech, AL; catalog #0100-01). Samples were imaged either on a Nikon Eclipse TE2000 microscope (Nikon, Tokyo, Japan) with an Opti-Grid structured illumination imaging system using Volocity software (PerkinElmer, Waltham, MA, USA) or on a Nikon A1 Inverted Confocal Microscope (Nikon, Tokyo, Japan). All primary antibodies used in the study are listed in Table S1.

### Cell counts and tubule area quantification

For all cell counts, 5-15 random tubules within each testis, from at least 3 different sections, were analyzed (n=3 testes from independent animals). The Fiji/ImageJ (NIH) Cell Counter plug-in was used to manually count positive cells per tubule. For measuring tubule cross-sectional areas, the perimeters of individual tubules were outlined using the Polygon Selection tool, and the area in pixels was calculated with the Measure function. Pixel area was then converted to square microns based on the pixel dimensions of each image. Graph results are shown as mean ± SD. Statistical analyses were performed using a two-tailed Student t-test, and a *P* value of P<0.05 was considered statistically significant.

### RNA extraction, cDNA synthesis, and quantitative real-time PCR (qPCR)

Total RNA was extracted and processed for qPCR. Tissue was homogenized by vortexing in 800µl TRIzol reagent (Invitrogen/Thermo Fisher). RNA extraction was then performed using a TRIzol/isopropanol precipitation method. Briefly, 200 µL of chloroform was added to the Trizol/tissue mixture, shaken by hand, incubated at room temperature for 3 minutes, and centrifuged at 12,000 x g for 10 minutes at 4°C. The upper aqueous layer was carefully recovered and added to 400 µL isopropanol, which was rocked at room temperature for 10 minutes. After centrifugation at 12,000 x g for 10 minutes at 4°C, supernatant was removed and the pellet was washed with 500 µL of ethanol. After another centrifugation (with same parameters), the RNA pellet was briefly air-dried and diluted in nuclease-free water. RNA quality was assessed by spectrophotometric analysis via absorbance at 260 and 280 nm, in which only RNA samples with a 260/280 ratio greater than or equal to 1.6 was used for qPCR analysis (although sample ratios were usually between 1.7-2.0). An iScript cDNA synthesis kit (BioRad) was used on 500ng of RNA for cDNA synthesis. qPCR was performed using the Fast SYBR Green Master Mix (Applied Biosystems/Thermo Fisher) on the StepOnePlus Real-Time PCR system (Applied Biosystems/Thermo Fisher). The following parameters were used: 95°C for 20s, followed by 40 cycles of 95°C for 3s and 60°C for 30s, followed by a melt curve run. Primer specificity for a single amplicon was verified by melt curve analysis. *Gapdh* was used as an internal normalization control. Fold change in mRNA levels was calculated relative to controls using a ΔΔCt method. A two-tailed Student t-test was performed to calculate *P* values based on delta C_t_ values, in which *P*<0.05 was considered statistically significant. Sequences of primers used in this study are listed in Table S2.

### Biotin tracer injections

P30 or P90 male mice (n=3 males for each age) were anesthetized via inhaled isoflurane before a small incision was made in the lower abdomen, exposing the abdominal cavity. Using forceps to grip the epididymal fat pads, the testis was then gently pulled out of the abdomen. Twenty microliters of either 1mM CaCl_2_ (in PBS) alone for one testis, or 1mM CaCl_2_ with 10 mg/mL Biotin (EZ-Link™ Sulfo-NHS-LC-Biotin, No-Weigh Format; Thermo Fisher #A39257) for the contralateral experimental testis was injected. Testes were then replaced into the abdominal cavity and the incision closed. Mice were kept under anesthesia via isoflurane for an additional 30 minutes before being euthanized by cervical dislocation. Testes were harvested, fixed, and prepared for cryosectioning and immunofluorescence as described above. To detect biotin binding in testicular cryosections, Alexa-Fluor-555-conjugated streptavidin (Thermo Fisher #S21381, 2 µg/ml) was used for 1 hour at room temperature, along with Hoechst 33342 dye to stain nuclei.

### Statistical Analysis

Statistical details of experiments, such as the exact value of n, what n represents, precision measures (mean ± SD), and statistical significance can be found in the Figure Legends. Statistical analysis was conducted using Excel (Microsoft). The results are reported as mean ± standard deviation. For cell counts and tubule area quantification, a two-tailed Student t-test was performed on control versus cKO samples. For qPCR, a two-tailed Student t-test was performed on delta C_t_ values (normalized to *Gapdh*) for each gene for control versus cKO samples. *P*<0.05 was considered statistically significant.

### Data and Code Availability

This study did not generate or analyze any unique datasets or code.

## Notes

### Competing Interest Statement

The authors have declared no competing interest.

