## Supplemental Data for "*Cdc42* activity in Sertoli cells is essential for maintenance of spermatogenesis"

### SUPPLEMENTAL INFORMATION FIGURE TITLES AND LEGENDS

**Figure S1. *Cdc42* is expressed in Sertoli cells throughout fetal development and during adulthood.**

(A) Plot showing *Cdc42* expression levels, which were generated from fetal gonad cell-type-specific microarray data (Jameson et al., 2012), where cell types (i.e., supporting Sertoli cells, interstitial cells, endothelial cells, and germ cells) were each plotted in different colors. Plot shows expression levels for each cell type in E11.5, E12.5, and E13.5 XY gonads. Expression values below 6 are generally considered background expression. *Cdc42* is expressed at high levels in all cell types throughout all 3 stages. (B-D) Violin plots showing single-cell RNA-Seq data for fetal (B and C) and adult (D) testes, from previously published datasets (Green et al., 2018; Stevant et al., 2018), showing expression plotted by either stage (B) or cell type (C and D). Plots were generated using The ReproGenomics Viewer (Darde et al., 2019; Darde et al., 2015).

**Figure S2. Adult cKO testes exhibit a complete loss of spermatogenic stages of germ cells.**

(A-F) Immunofluorescence images of P90 adult control *Dhh*-Cre;*Cdc42*$^{flox/+}$ (A,C,E) and cKO (*Dhh*-Cre;*Cdc42*$^{flox/flox}$) (B,D,F) testes. A'-F' are higher-magnification images of the boxed regions in A-F. Dashed lines indicate tubule boundaries. (A and B) GFRA1+ undifferentiated spermatogonia are located basally within control (A) tubules (arrowhead in A'), but are absent in cKO (B) tubules. (C and D) Whereas control (C) tubules contain ZBTB16+ undifferentiated spermatogonia (arrowhead in C') and STRA8+ preleptotene spermatocytes (arrow in C'), these cell types cannot be detected in cKO (D) tubules. (E and F) While control (E) tubules contain H1T-positive pachytene spermatocytes with XY-body-enriched γH2AX staining (arrows in E') and H1T-positive round spermatids (arrowheads in E'), cKO (F) tubules do not. Scale bars, 100 µm. (G) qPCR analyses of germ-cell-specific gene expression in P60 adult cKO testes relative to control *Dhh*-Cre;*Cdc42*$^{flox/+}$ testes (n=4 testes each for controls and cKO, all from independent males). Data in G is shown as mean fold change ± SD. *P* values were calculated via a two-tailed Student t-test.

**Figure S3. Onset of quiescence in postnatal Sertoli cells occurs normally in cKO testes.**

(A-D) Immunofluorescence images of P15 (A and B) and P24 (C and D) control *Dhh*-Cre;*Cdc42*$^{flox/+}$ (A and C) and cKO (*Dhh*-Cre;*Cdc42*$^{flox/flox}$) (B and D) testes. A'-D', B", and D" are higher-magnification images of the boxed regions in A-D. (A-D) P15 (A and B) and P24 (B and D) Sertoli cells in control (A and C) and cKO (B and D) testes are MKI67-negative. Rare MKI67+ Sertoli cells are observed in both control and cKO testes (arrow in D'). Even Sertoli cell nuclei mislocalized in the middle of cKO tubules at P15 (arrowheads in B") or P24 (arrowheads in D") are MKI67-negative. Scale bars, 50 µm.

**Figure S4. Sustained BTB disruption in cKO testes is not due to defects in the interstitial compartment.**

(A-L) Immunofluorescence images of P90 control *Dhh*-Cre;*Cdc42*$^{flox/+}$ (A,C,E,G,I,K) and cKO (*Dhh*-Cre;*Cdc42*$^{flox/flox}$) (B,D,F,H,J,L) testes. A',B',G', and H' are higher-magnification images of the boxed regions in A,B,G, and H. (A and B) Biotin tracer injection reveals only limited penetration of biotin into control (A) tubules, but extensive presence of biotin within the center of cKO (B) tubules. (C and D) Compared to controls (C), cKO (D) testes contain similarly abundant CYP17A1+ Leydig cells in the interstitium. (E and F) Both control (E) and cKO (F) testes display extensive vascularization (PECAM1+ cells). (G and H) Relative to control (G) testes, cKO (H) testes show increased cell cycle activity (MKI67 immunoreactivity) in interstitial cells such as peritubular myoid cells (arrows in G' and H'). Arrowhead in H denotes MKI67+ cKO Sertoli cell. (I and J) While control (I) testes have CD45+ immune cells such as macrophages (AIF1+) in the interstitium, cKO (J) testes exhibit increased CD45-bright immune cell infiltration (likely T cells; arrows in L) in the interstitium, as well as ectopic macrophage presence within tubule lumens (arrowheads in L). Scale bars, 100 µm. (M) qPCR analyses of interstitial gene expression in P60 adult cKO testes relative to control *Dhh*-Cre;*Cdc42*$^{flox/+}$ testes (n=4 testes each for controls and cKO, all from independent males). *Cyp11a1* and *Cyp17a1* are specific to Leydig cells; *Kit* is expressed in Leydig cells, as well as differentiating spermatogonia in controls; and *Cdh5* is specific to vascular endothelial cells. Data in M is shown as mean fold change ± SD. *P* values were calculated via a two-tailed Student t-test. ns, not significant (*P*>0.05).

**Figure S5. Early postnatal differentiation of Sertoli and germ cells occurs normally in cKO testes.**

(A-H) Immunofluorescence images of P7 control *Dhh*-Cre;*Cdc42*$^{flox/+}$ (A,C,E,G) and cKO (*Dhh*-Cre;*Cdc42*$^{flox/flox}$) (B,D,F,H) testes. G'a, G'b, H'a, and H'b are higher-magnification images of the boxed regions in G and H. (A and B) Compared to controls (A), cKO testes have similar numbers of Sertoli (GATA4+) and germ (TRA98+) cells. (C and D) GATA1 expression is detected in both control (C) and cKO (D) Sertoli cells. (E and F)

Both AMH and AR are detected within control (E) and cKO (F) testes, but AMH expression seems slightly decreased in cKO Sertoli cells. (G and H) Similar levels of CC3+ apoptotic cells are observed in control (G) versus cKO (H) testes; co-stains indicate most apoptotic cells are germ cells (arrowheads in G'a, G'b', H'a, and H'b) with Sertoli cells only rarely detected undergoing cell death (arrows in G'b' and H'b). Scale bars, 100 µm.

**Figure S6. Early postnatal differentiation of interstitial cells occurs normally in cKO testes.**

(A-F) Immunofluorescence images of P7 control *Dhh*-Cre;*Cdc42*$^{flox/+}$ (A,C,E) and cKO (*Dhh*-Cre;*Cdc42*$^{flox/flox}$) (B,D,F) testes. (A and B) Both control (A) and cKO (B) testes exhibit extensive interstitial vascularization by PECAM1+ endothelial cells. (C and D) Similar to controls (C), cKO (D) testes contain CYP17A1+ Leydig cells. (E and F) No gross differences are observed in overall cell number of CD45+ immune cells in control (E) versus cKO (F) testes. Scale bars, 100 µm.

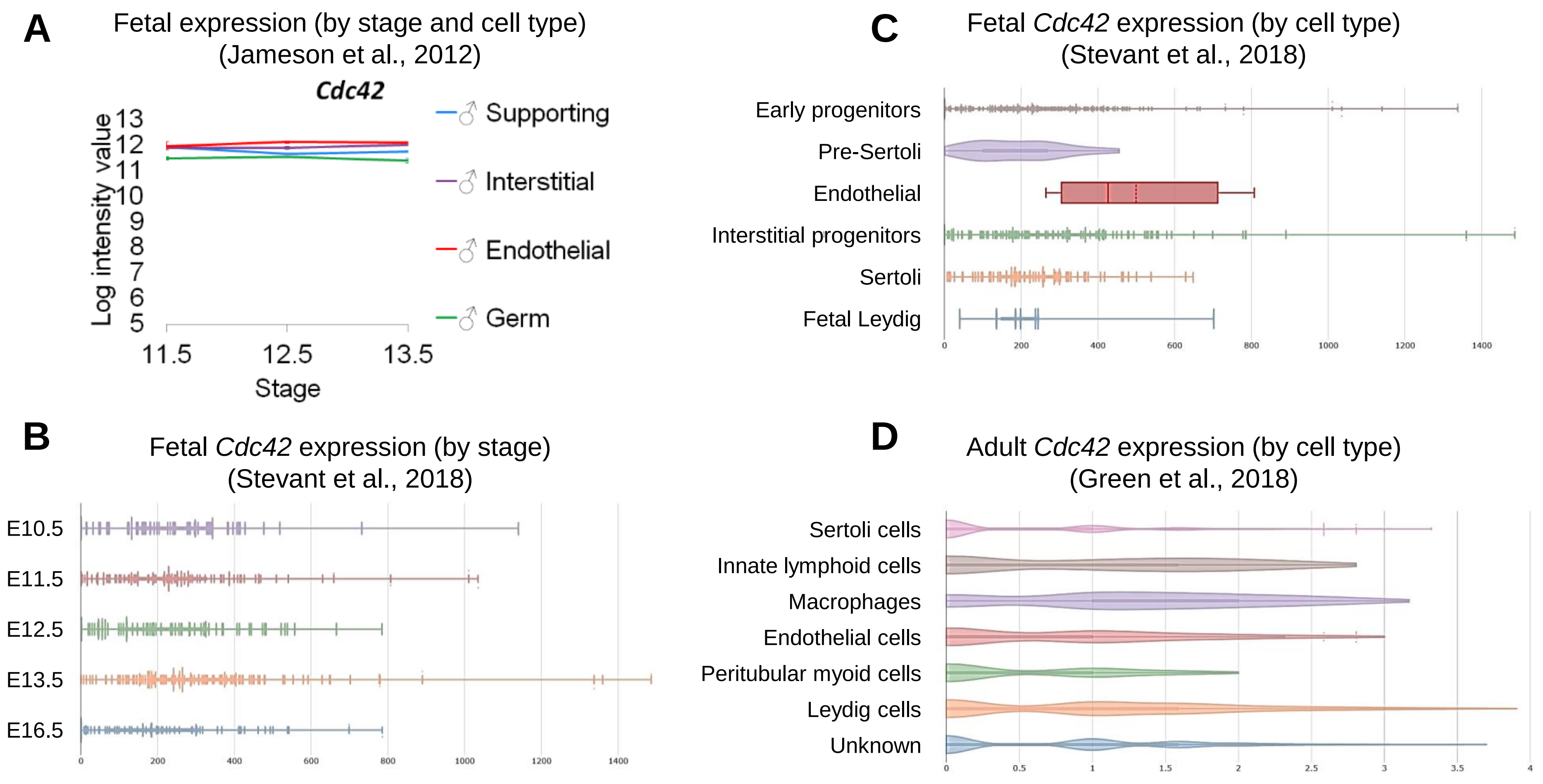
A
Fetal expression (by stage and cell type)
(Jameson et al., 2012)
Cdc42
Log intensity value
13
12
11
10
9
8
7
6
5
11.5
12.5
13.5
Stage
♂ Supporting
♂ Interstitial
♂ Endothelial
♂ Germ
B
Fetal Cdc42 expression (by stage)
(Stevant et al., 2018)
E10.5
E11.5
E12.5
E13.5
E16.5
0
200
400
600
800
1000
1200
1400
C
Fetal Cdc42 expression (by cell type)
(Stevant et al., 2018)
Early progenitors
Pre-Sertoli
Endothelial
Interstitial progenitors
Sertoli
Fetal Leydig
0
200
400
600
800
1000
1200
1400
D
Adult Cdc42 expression (by cell type)
(Green et al., 2018)
Sertoli cells
Innate lymphoid cells
Macrophages
Endothelial cells
Peritubular myoid cells
Leydig cells
Unknown
0
0.5
1
1.5
2
2.5
3
3.5
4

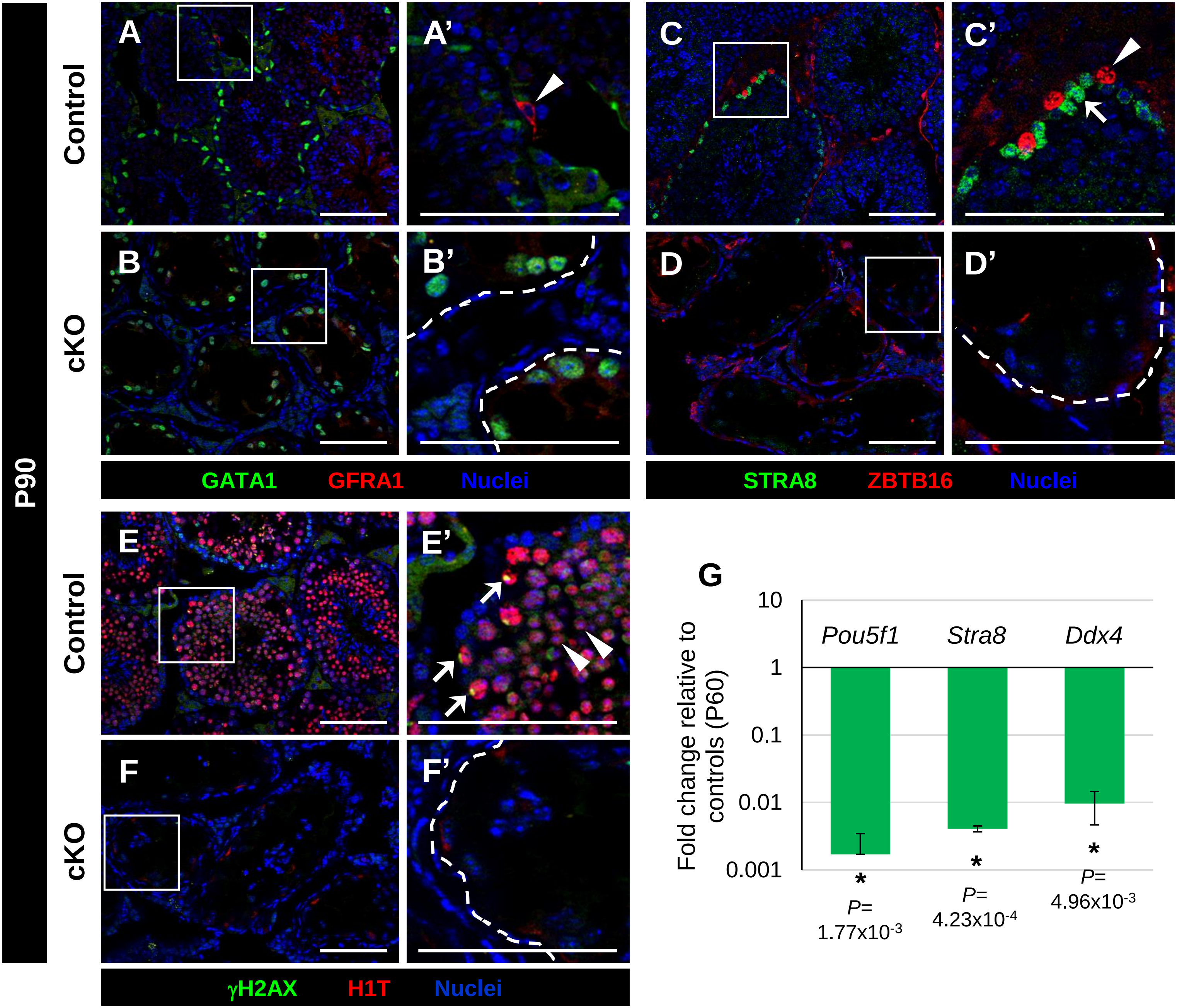
P90
Control
cKO
Control
cKO
A
A'
B
B'
C
C'
D
D'
E
E'
F
F'
GATA1
GFRA1
Nuclei
STRA8
ZBTB16
Nuclei
γH2AX
H1T
Nuclei
G
Fold change relative to controls (P60)
10
1
0.1
0.01
0.001
Pou5f1
Stra8
Ddx4
*
P=
1.77x10-3
*
P=
4.23x10-4
*
P=
4.96x10-3

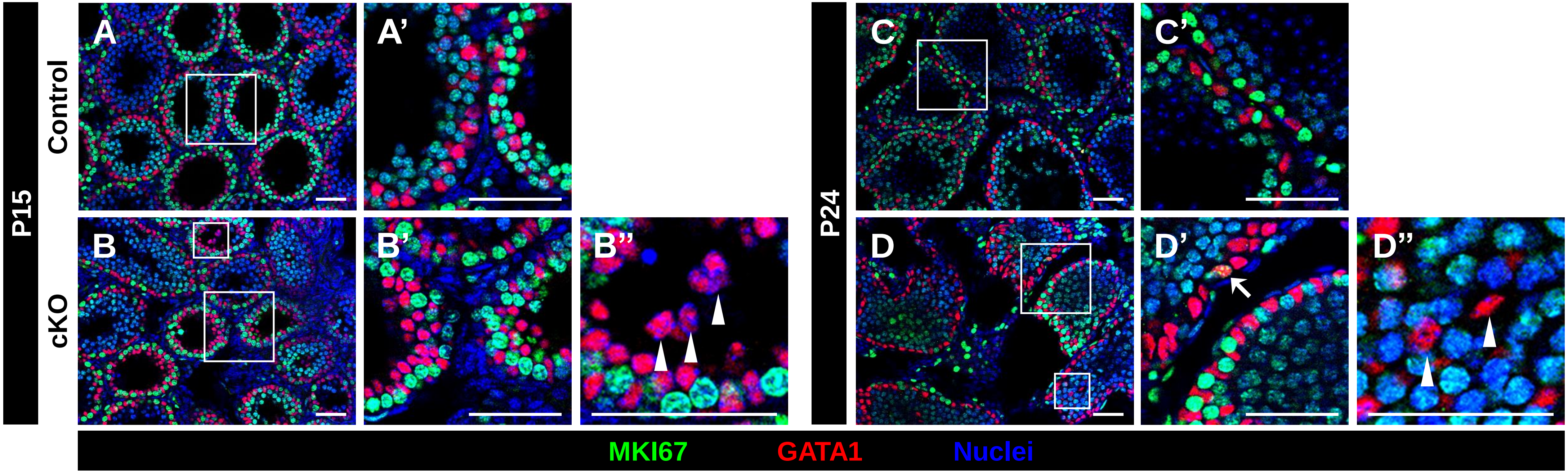
P15
Control
cKO
A
A'
B
B'
B''
P24
C
C'
D
D'
D''
MKI67
GATA1
Nuclei

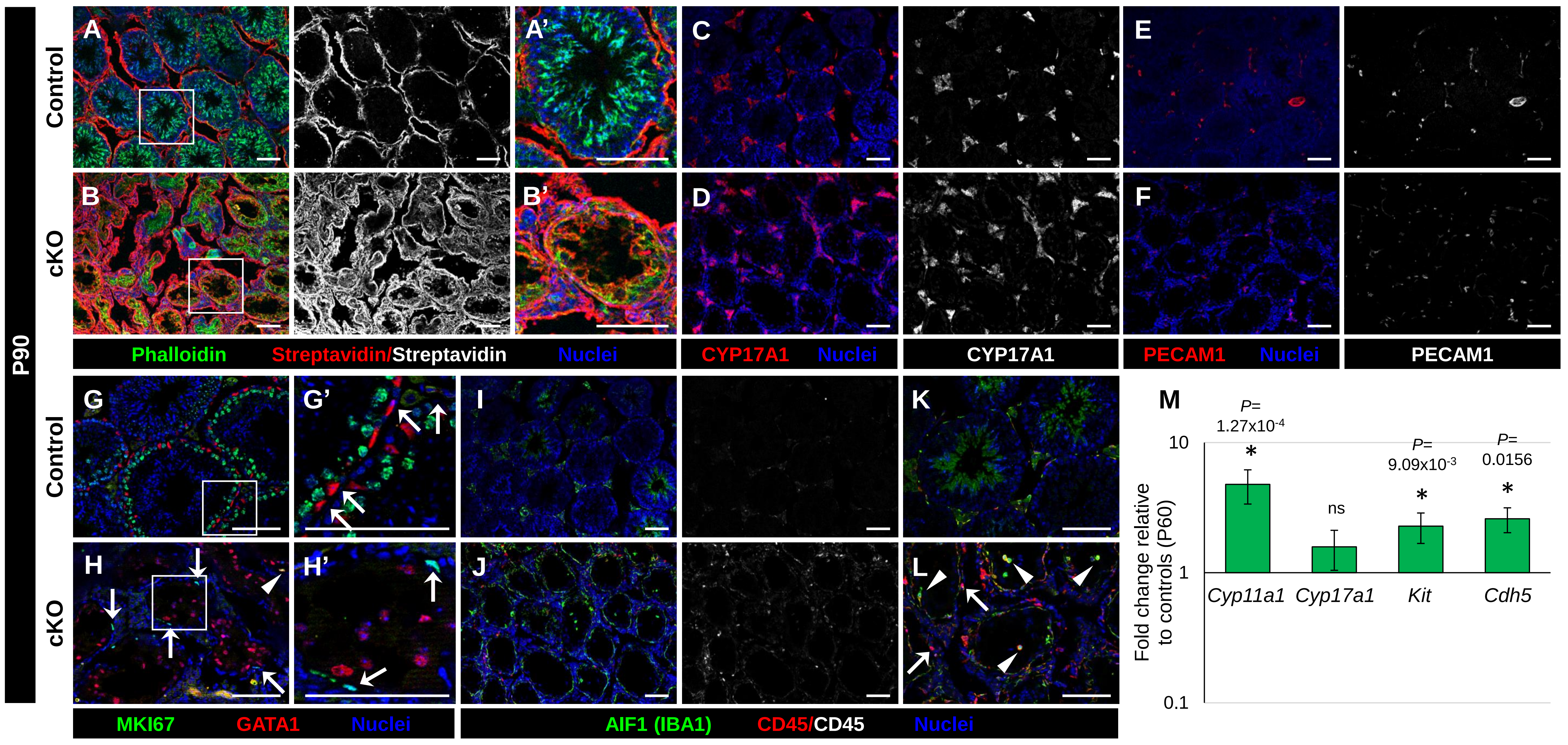
P90
Control
cKO
A
A'
B
B'
C
D
E
F
Phalloidin
Streptavidin/Streptavidin
Nuclei
CYP17A1
Nuclei
CYP17A1
PECAM1
Nuclei
PECAM1
G
G'
H
H'
I
J
K
L
MKI67
GATA1
Nuclei
AIF1 (IBA1)
CD45/CD45
Nuclei
M
P=
1.27x10-4
P=
9.09x10-3
P=
0.0156
ns
10
1
0.1
Fold change relative
to controls (P60)
Cyp11a1
Cyp17a1
Kit
Cdh5

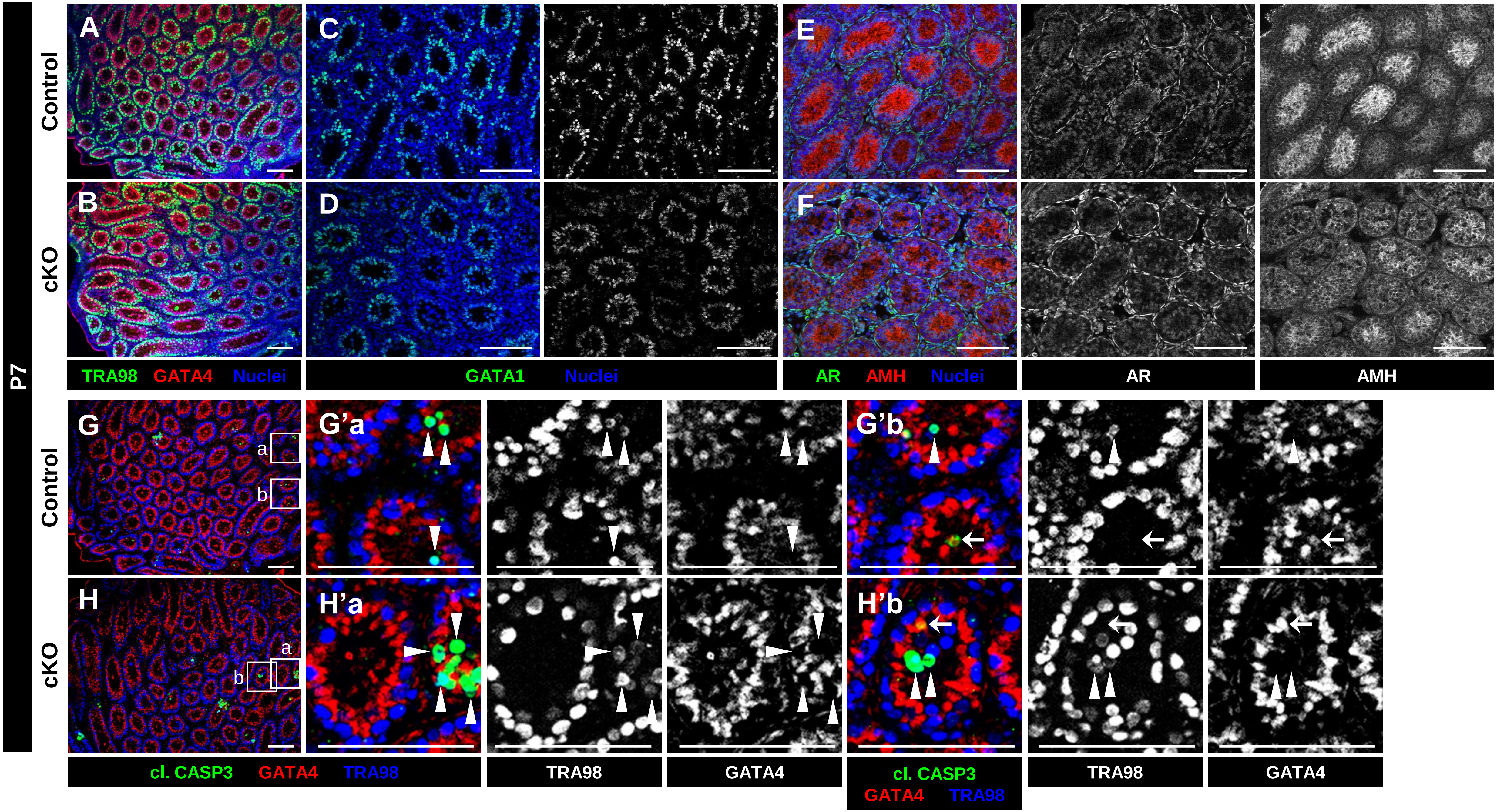

P7
Control
cKO
A
B
C
D
E
F
TRA98 GATA4 Nuclei
GATA1 Nuclei
AR AMH Nuclei
AR
AMH
G
H
a
b
G'a
G'b
H'a
H'b
cl. CASP3 GATA4 TRA98
TRA98
GATA4
cl. CASP3 GATA4 TRA98
TRA98
GATA4

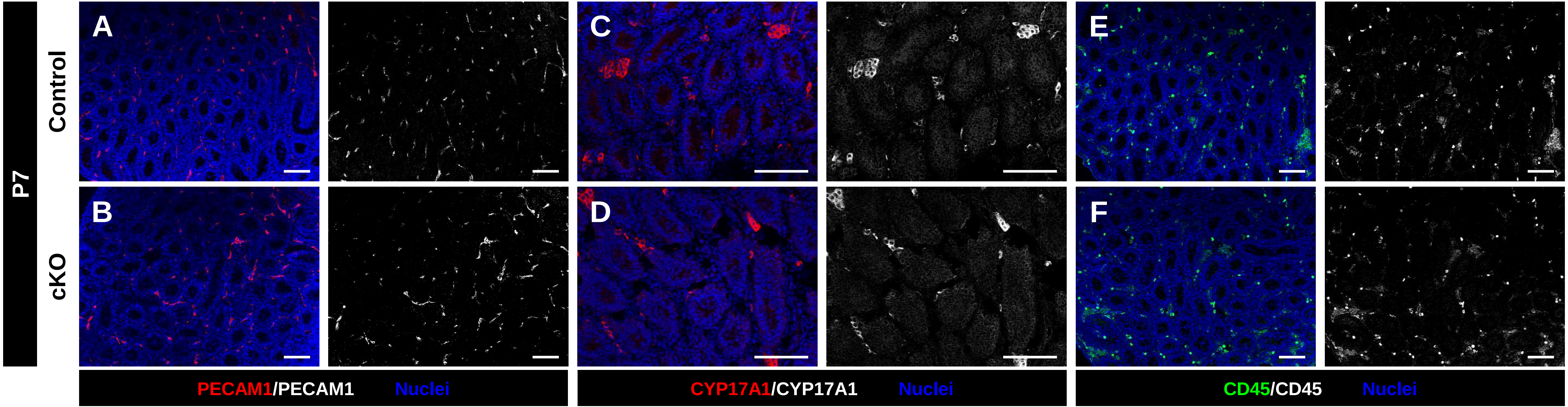
P7
Control
cKO
A
B
C
D
E
F
PECAM1/PECAM1 Nuclei
CYP17A1/CYP17A1 Nuclei
CD45/CD45 Nuclei

**Table S1. Primary antibodies.** List of primary antibodies used for immunofluorescence in this study (in alphabetical order).

| Primary Antibody | Dilution | Company/Source |
|---|---|---|
| Rabbit anti-AIF1 (IBA1) | 1:1,000 | Wako #019-19741 |
| Goat anti-AMH (MIS) | 1:500 | Santa Cruz #sc-6886 |
| Rabbit anti-AR | 1:300 | Santa Cruz #sc-816 |
| Rat anti-CD45 | 1:300 | BioLegend #103101 |
| Goat anti-KIT (C-KIT) | 1:400 | R&D #AF1356 |
| Rabbit anti-Cleaved Caspase 3 (Asp175) | 1:250 | Cell Signaling #9661S |
| Rat anti-CDH1 | 1:500 | Thermo Fisher #13-1900 |
| Rabbit anti-CLDN11 | 1:1,000 | Thermo Fisher #36-4500 |
| Goat anti-CTNNB1 | 1:400 | Santa Cruz #sc-1496 |
| Goat anti-CYP17A1 | 1:500 | Santa Cruz #sc-46081 |
| Rabbit anti-DDX4 (MVH) | 1:1,000 | Abcam #ab13840 |
| Rat anti-GATA1 | 1:1,000 | Santa Cruz #sc-265 |
| Goat anti-GATA4 | 1:100 | Santa Cruz #sc-1237 |
| Rabbit anti-GDNF | 1:500 | Santa Cruz #sc-328 |
| Rabbit anti-γH2AX | 1:1,000 | Millipore-Sigma #07-164 |
| Guinea pig anti-H1T | 1:2,000 | Mary Ann Handel |
| Goat anti-ITGB1 | 1:500 | Novus #AF2405-SP |
| Rabbit anti-MKI67 (Ki67) | 1:300 | Thermo Fisher #89351-224 |
| Rabbit anti-PARD3 (Par3) | 1:100 | Novus #NBP1-88861 |
| Goat anti-PECAM1 | 1:300 | R&D #AF3628 |
| Rat anti-PECAM1 | 1:250 | BD Biosciences #553370 |
| Alexa 647 Phalloidin (in methanol) | 1:500 | Thermo Fisher #A22287 |
| Rhodamine Phalloidin (in methanol) | 1:500 | Thermo Fisher #R415 |
| Rabbit anti-SCRIB | 1:300 | Santa Cruz #sc-28737 |
| Rabbit anti-Phospho-SCRIB | 1:500 | Cell Signaling ##12316 |
| Rabbit anti-SOX9 | 1:4,000 | Millipore-Sigma #AB5535 |
| Rabbit anti-STRA8 | 1:3,000 | Abcam #ab49602 |
| Rat anti-TRA98 | 1:1,000 | Abcam #ab82527 |
| Mouse anti-TUBB3 (TUJ1) | 1:1,000 | BioLegend #801201 |
| Chicken anti-Vimentin | 1:1,000 | BioLegend #919101 |
| Rat anti-Vimentin | 1:1,000 | BioLegend #699301 |
| Mouse anti-ZBTB16 (PLZF) | 1:250 | Millipore #OP128-100UG |

**Table S2. qPCR primers.** Sequences of primers used for quantitative real-time PCR (qPCR) in this study (in alphabetical order).

| Gene name | Sequence (5' to 3') |
|---|---|
| *Amh* forward | CCACACCTCTCTCCACTGGTA |
| *Amh* reverse | GGCACAAAGGTTCAGGGGG |
| *Ar* forward | CAGGAGGTAATCTCCGAAGGC |
| *Ar* reverse | ACAGACACTGCTTTACACAACTC |
| *Cdh5* forward | TCCTCTGCATCCTCACTATCACA |
| *Cdh5* reverse | GTAAGTGACCAACTGCTCGTGAAT |
| *Cldn11* forward | ATGGTAGCCACTTGCCTTCAG |
| *Cldn11* reverse | AGTTCGTCCATTTTTCGGCAG |
| *Cyp11a1* forward | GGAGGAAGCCGACAACAATGA |
| *Cyp11a1* reverse | TCCACCTCACACGGTTCTCAA |
| *Cyp17a1* forward | CAGAGAAGTGCTCGTGAAGAAG |
| *Cyp17a1* reverse | AGGAGCTACTACTATCCGCAAA |
| *Ddx4* forward | TACTGTCAGACGCTCAACAGGA |
| *Ddx4* reverse | ATTCAACGTGTGCTTGCCCT |
| *Fshr* forward | GGGATCTGGATGTCATCACT |
| *Fshr* reverse | GGAGAACACATCTGCCTCTA |
| *Gapdh* forward | AGGTCGGTGTGAACGGATTTG |
| *Gapdh* reverse | TGTAGACCATGTAGTTGAGGTCA |
| *Gata1* forward | TGGGGACCTCAGAACCCTTG |
| *Gata1* reverse | GGCTGCATTTGGGGAAGTG |
| *Gdnf* forward | GACTTGGGTTTGGGCTATGA |
| *Gdnf* reverse | AACATGCCTGGCCTACTTTG |
| *Inhbb* forward | GAGCGCGTCTCCGAGATCATCA |
| *Inhbb* reverse | CGTACCTTCCTCCTGCTGCCCTT |
| *Kit* forward | CATGGCGTTCCTCGCCT |
| *Kit* reverse | GCCCGAAATCGCAAATCTTT |
| *Pdgfa* forward | GACGGTCATTTACGAGATACCTC |
| *Pdgfa* reverse | CTACGCCTTCCTGTCTCCTC |
| *Pou5f1* forward | GGAGGAAGCCGACAACAATGA |
| *Pou5f1* reverse | TCCACCTCACACGGTTCTCAA |
| *Sox9* forward | GCGGAGCTCAGCAAGACTCTG |
| *Sox9* reverse | ATCGGGGTGGTCTTTCTTGTG |
| *Stra8* forward | GCAGGTTGAAGGATGCTTTGAGC |
| *Stra8* reverse | CCTAAGGAAGGCAGTTTACTCCCAGTC |
